# Phylogenomic analyses in the apple genus *Malus* s.l. reveal widespread hybridization and allopolyploidy driving the diversifications, with insights into the complex biogeographic history in the Northern Hemisphere

**DOI:** 10.1101/2021.10.12.464085

**Authors:** Bin-Bin Liu, Chen Ren, Myounghai Kwak, Richard G.J. Hodel, Chao Xu, Jian He, Wen-Bin Zhou, Chien-Hsun Huang, Hong Ma, Guan-Ze Qian, De-Yuan Hong, Jun Wen

## Abstract

Phylogenomic evidence from an increasing number of studies has demonstrated that different data sets and analytical approaches often reconstruct strongly supported but conflicting relationships. In this study, hundreds of single-copy nuclear (SCN) genes (785) and complete plastomes (75) were used to infer the phylogenetic relationships and estimate the historical biogeography of the apple genus *Malus* sensu lato, an economically important lineage disjunctly distributed in the Northern Hemisphere involved in known and suspected hybridization and allopolyploidy events. The nuclear phylogeny recovered the monophyly of *Malus* s.l. (including *Docynia*); however, it was supported to be biphyletic in the plastid phylogeny. An ancient chloroplast capture event best explains the cytonuclear discordance that occurred in the Eocene in western North America. Our conflict analysis demonstrated that ILS, hybridization, and allopolyploidy could explain the widespread nuclear gene tree discordance. We detected one deep hybridization event (*Malus doumeri*) involving the ancestor of pome-bearing species and *Docynia delavayi*, and one recent hybridization event (*Malus coronaria*) between *M. sieversii* and a combined clade of *M. ioensis* and *M. angustifolia*. Furthermore, our historical biogeographic analysis combining living and fossil species supported a widespread East Asian-western North American origin of *Malus* s.l., followed by a series of extinction events in the Eocene in northern East Aisa and western North America. This study provides a valuable evolutionary framework for the breeding and crop improvement of apples and their close relatives.

## Introduction

The apple genus *Malus* Mill. (tribe Maleae, Rosaceae) is of great economic importance, with the domesticated apple (*M. domestica* (Suckow) Borkh.), various crabapples, as well as some important ornamentals (e.g., *M. halliana* Koehne, *M. hupehensis* (Pamp.) Rehder, and *M. micromalus* Makino)^1^. *Malus* comprises ca. 38-55 species disjunctly distributed across the temperate Northern Hemisphere^2, 3^, from East Asia (ca. 27 species)^2^, Central Asia (ca. two species), Europe (three species)^4^, and the Mediterranean (ca. two species)^4^ to western (one species) and eastern North America (three species)^3^. Hybridization, polyploidy, and apomixis within the apple genus *Malus* have been reported to have occurred frequently in the wild and horticulture, and introgression has led to many taxa described in *Malus*^3, 5^. These reticulated processes have significantly challenged the accurate phylogenetic inference of *Malus*; however, this genus also provides an ideal case for studying the phylogenomic discordance and its underlying potential causes. Based on morphology, many taxonomists have struggled to propose a reasonable classification system^5–15^, and over-emphasis of one or a few morphological characters resulted in the controversial taxonomy at the sectional level, varying from two, five, six, to eight sections^15–17^. It has been challenging to elucidate the evolutionary relationships among *Malus* member*s* based solely on morphological evidence. Robinson et al. (2001) first used molecular evidence (plastid *matK* and nuclear ribosomal internal transcribed spacer (nrITS) sequences) to explore the phylogenetic relationships of five sections sensu Rehder (1940). Subsequently, phylogenetic studies on *Malus* and its close relatives used more plastid and nrITS sequences^18^, whole plastome and/or entire nuclear ribosomal DNA (nrDNA)^19–21^, and transcriptomic dataset^22^. These studies provided significant insights into the major clades, but many questions remain due to the limited informative sites and/or taxon sampling (Fig. 1). The phylogenetic analysis in the framework of Maleae so far has shown strongly supported yet discordant nuclear and chloroplast topologies of *Malus* sect. *Chloromeles* (Decne.) Rehder and *M.* sect. *Eriolobus* lineage, which makes *Malus* s.l. either monophyletic (nuclear sequences: Fig. 1f-i) or biphyletic (whole plastome: Fig. 1e), suggesting possible hybridization and/or chloroplast capture events in the origin of this clade^21^. However, Lo & Donoghue (2012) produced a monophyletic *Malus* s.l. based on 11 plastid regions as Robinson et al. (2001) did using *matK* regions (Fig. 1a, b). Generally, phylogenetic relationships within *Malus* s.l. are not fully resolved, particularly along the backbone of the phylogeny.

**Fig. 1.**
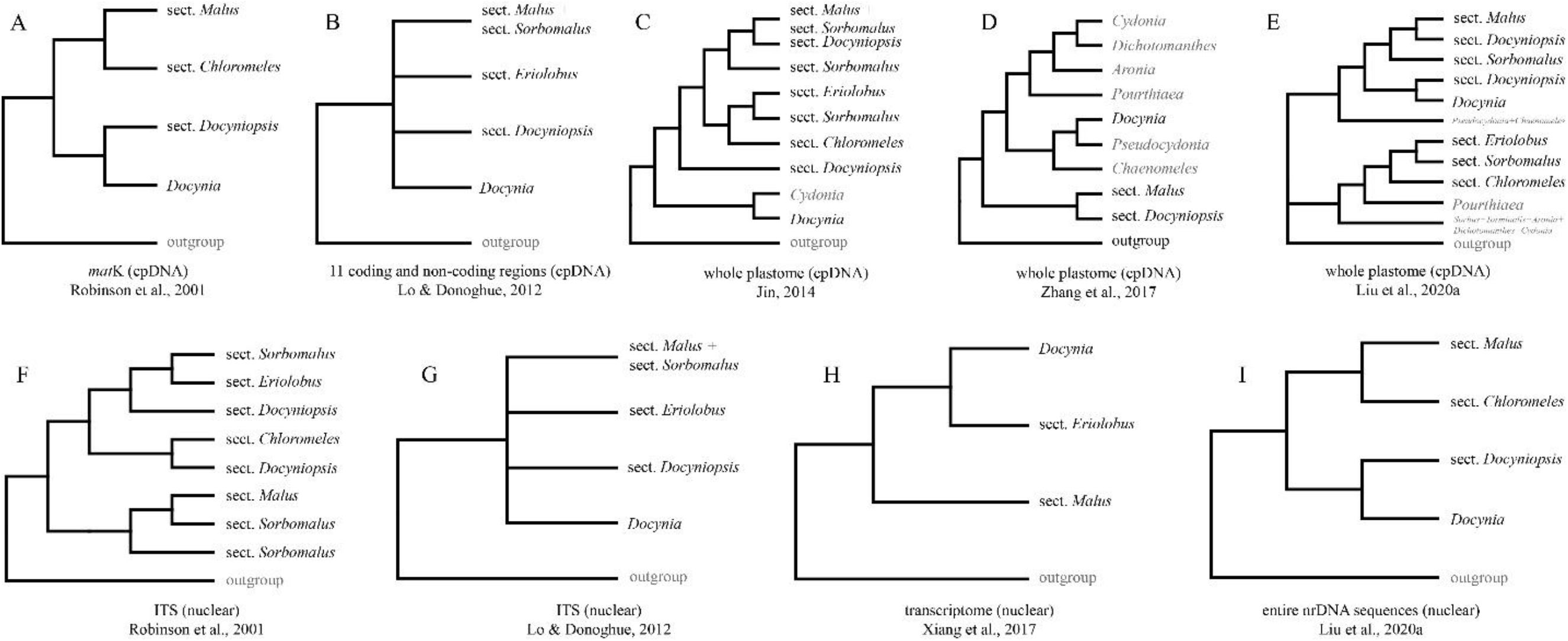
Phylogenetic hypotheses among sections of *Malus* s.l. estimated by previous studies. **a**, plastid *matK* sequence^17^; **b**, 11 plastid coding and non-coding regions^18^; **c**, whole plastome^19^; **d**, whole plastome^20^; **e**, whole plastome^21^; **f**, nuclear ITS sequence^17^; **g**, nuclear ITS sequence^18^; **h**, transcriptome^22^; **i**, entire nrDNA sequences^21^. The sectional delimitation in *Malus* s.l. followed Phipps et al. (1990).

Phylogenomics has revolutionized plant systematics and evolution as well as the associated fields in the last decades, enabling the utilization of a large number of nuclear genes for testing phylogenetic hypotheses, especially untangling recalcitrant relationships using traditional molecular systematic approaches^23–25^. While data collection efforts have significantly increased overall phylogenetic support, these analyses are a double-edged sword by demonstrating that different data sets and analytical approaches often reconstruct strongly supported but conflicting relationships^26–28^. Underlying these conflicting topologies are strongly discordant gene trees^29^, which may be due to several biological processes such as gene duplication, incomplete lineage sorting (ILS), and gene flow (e.g., hybridization, allopolyploidy, and introgression). High levels of gene tree discordance have occurred in the nuclear genome and in the plastome, with the latter often regarded as a single locus^30–34^. The evidence so far suggests that gene flow between lineages promoted the diversification of land plants^34, 35^. Additionally, conflicts between nuclear and organellar genomes, often called cytonuclear discordance, complicate phylogenetic inference. Generally, cytonuclear conflict has been interpreted as a result of introgressive chloroplast capture, which has been found in various lineages of angiosperms^21, 36–38^. Despite widespread introgression detected in angiosperms, current methods for phylogenetic inference often either assume no gene tree discordance (i.e., concatenated supermatrices) or only consider a coalescent model in which all discordance is attributed to ILS^39, 40^. Recently, through calculating unique, conflicting, and concordant bipartitions (e.g., *phyparts*)^29^ or quartet-based evaluation (e.g., Quartet-Sampling)^41^, the highly supported but sometimes conflicting relationships can be quantified. Additionally, phylogenetic network analysis has allowed estimating species trees to account for ILS and introgression^42–44^, and this approach can explore the mechanisms of the discordance.

*Malus* s.l. has a wide disjunct distribution in the major refugia of the Northern Hemisphere: East Asia, the Mediterranean, western North America, and eastern North America, as well as Europe. This distribution pattern provides an ideal case study for exploring the evolution of the major patterns of biogeographic disjunctions in the Northern Hemisphere. Such disjunctions are generally considered to be remnants of a more continuously distributed, mixed mesophytic forest during the Tertiary^45, 46^. Due to the geologic and climatic oscillations, the once more widely distributed flora was fragmented and became relict in four major refugia: East Asia, eastern North America, western North America, and Southwest Europe. Based on phylogenetic analyses of 47 chloroplast genomes, Nikiforova et al. (2013) suggested a North American origin of *Malus*, which was in accordance with the fossil evidence, because many fossils were recorded in the middle to late Eocene from western North America^47–53^. Contrasting this New World origin hypothesis, Jin (2014) proposed an alternative hypothesis of East Asian origin based on biogeographic analyses using complete plastome data. Jin (2014) concluded that this conflict may be due to the lack of sampling of key early diverged lineages, such as *Malus doumeri* (Bois) A. Chev., *M. florentina* C.K.Schneid., and *M. trilobata* C.K.Schneid. in Nikiforova et al. (2013)’s study. However, Jin (2014)’s misidentified *M. doumeri* sample (actually *Pseudocydonia sinensis* (Thouin) C.K.Schneid.^21^) may have also biased the phylogenetic inference and historical biogeographic estimation. These two conflicting hypotheses showcase the need for a robust phylogenetic framework with a comprehensive taxon sampling scheme to reconstruct the biogeographic history of *Malus*.

In this study, we intend to explore gene tree concordance and the phylogenetic relationships among major clades to test the potential ILS and hybridization events in the evolutionary history of *Malus* s.l. We further conduct biogeographic analyses to infer geographic origins and timing of possible hybrid clades. Due to their significant economic importance, genomes of apples and their close relatives have been sequenced, such as *Malus baccata* (L.) Borkh.^54^, *M. × domestica*^55–59^, *M. sieversii* (Ledeb.) M.Roem.^59^, and *M. sylvestris* Mill.^59^, which provided substantial genome resources for exploring single-copy nuclear (SCN) genes for phylogenetic analysis. Additionally, numerous genome resequencing, transcriptomes, and raw genome skimming data are available from the NCBI sequence read archive (SRA: https://www.ncbi.nlm.nih.gov/sra). These raw genomic data, coupled with deep genome skimming data sensu Liu et al. (2021) generated for this study, provide an excellent opportunity to explore a robust phylogenetic framework within *Malus* s.l. using a large-scale phylogenomic analysis.

This study employs extensive genomic data, sampling from 77 individuals by integrating genome resequencing^57^, transcriptomic^22^, and deep genome skimming data^60^ to assemble 797 SCN genes and whole plastomes. Specifically, we aim to: (1) reconstruct a robust phylogenetic backbone of the apple genus *Malus* s.l.; (2) explore gene tree conflicts and evaluate the potential causes; and (3) investigate broad-scale biogeographic relationships and ancestral range evolution.

## Results

### Single-copy nuclear genes and plastome assembly

Raw reads for the 27 newly sequenced deep genome skimming data are available from the NCBI Sequence Read Archive (SRA: BioProject: PRJNA759205 with the accession for each sample listed in Supplementary Table S1). The number of clean reads ranged from 33 (*Cotoneaster salicifolius* var. *henryanus* (C.K.Schneid.) T.T.Yu) to 103 (*Pourthiaea zhejiangensis* (P.L.Chiu) Iketani & H.Ohashi) million with the sequencing depths varying from 6.7× to 20.8×, assuming an estimated genome size of around 750 Mb based on *Malus domestica* genome^55^.

We designed a set of 797 SCN genes from six genomes as mentioned below for this phylogenomic study on *Malus* and its close relatives. The number of genes recovered for each sample varied from 665 (83.4%: *Aronia melanocarpa* (Michx.) Elliott) to 797 (100%: 11 samples listed in Supplementary Table S2) (also referring to Fig. 2), and the number of genes after cleaning ranged from 568 (71.3%: *Malus doumeri*) to 785 (98.5%: *Malus baccata*) (Fig. 1 and Supplementary Table S2).

**Fig. 2.**
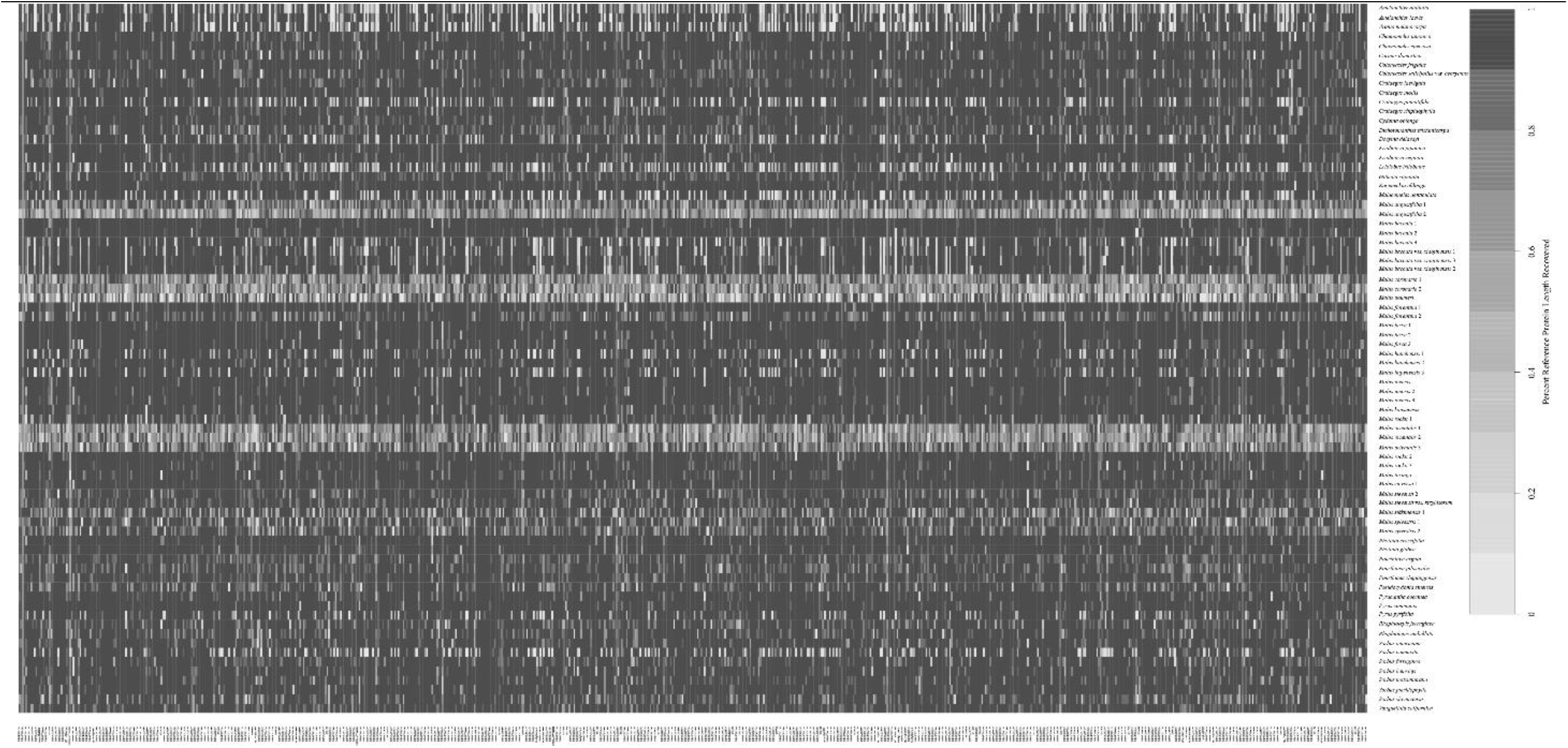
Heat map showing recovery efficiency for 797 genes recovered by HybPiper. Each column is a gene, and each row is one sample. The shade of gray in the cell is determined by the length of sequence recovered by the pipeline, divided by the length of the reference gene (maximum of 1.0).

We successfully assembled 69 plastomes for this study, except *Aronia melanocarpa*, *Crataegus pinnatifida* Bunge, *Eriolobus trilobatus* (Labill. ex Poir.) M.Roem., *Pyrus pyrifolia* (Burm.f.) Nakai, and *Sorbus commixta* Hedl. due to the limited plastid reads in the raw data. All plastomes were submitted to GenBank with accessions listed in Supplementary Table S1, and the aligned plastid matrix was deposited in Dryad Digital Repository (Data S4) https://doi.org/10.5061/dryad.2jm63xsq5.

### Nuclear phylogenetic analysis and gene tree discordance

We obtained sequences of 785 genes with at least 47 samples and 273 bp for each gene (Data S1, available from Dryad Digital Repository https://doi.org/10.5061/dryad.2jm63xsq5). We kept 604 genes with more than 900 bp in aligned length for the downstream phylogenetic analysis. To test the effect of missing data for phylogenetic analysis, we generated three datasets including different numbers of samples for each clean gene, i.e., 50%-sample dataset (604 genes), 80%-sample dataset (589 genes), and all-sample dataset (66 genes), and all these three datasets have been deposited in Dryad Digital Repository (Data S2-S4) https://doi.org/10.5061/dryad.2jm63xsq5. These three concatenated matrices consisted of 1,193,313 bp, 1,042,020 bp, and 152,891 bp in aligned length. We estimated nine phylogenetic trees (Fig. 3 and Supplementary Figs. S1-S9), in which six of them resulted in the same topology. Henceforth, we used the RAxML tree (Fig. 3: hereafter, referred to as “nuclear phylogeny”) estimated from the 80%-sample dataset for the following analysis. The nuclear phylogeny recovered *Malus* as paraphyletic with the genus *Docynia* Decne. embedded in *Malus*. *Malus* s.l. was delimited into three strongly supported major clades (BS = 100, 100, 100). Clade I included most of the species of *Malus* sect. *Malus* and *M.* sect. *Sorbomalus* except for a Mediterranean species, *M. florentina*. We sampled 27 individuals of clade I representing 11 species, disjunctly distributed between East Asia, Europe & Central Asia, and western North America. Clade II is composed of all the eastern North American (three species) and the Mediterranean species (two species). Clade III included two species, *M. doumeri*, previously delimited in *M.* sect. *Docyniopsis* C.K.Schneid. and *Docynia delavayi* C.K.Schneid.

**Fig. 3.**
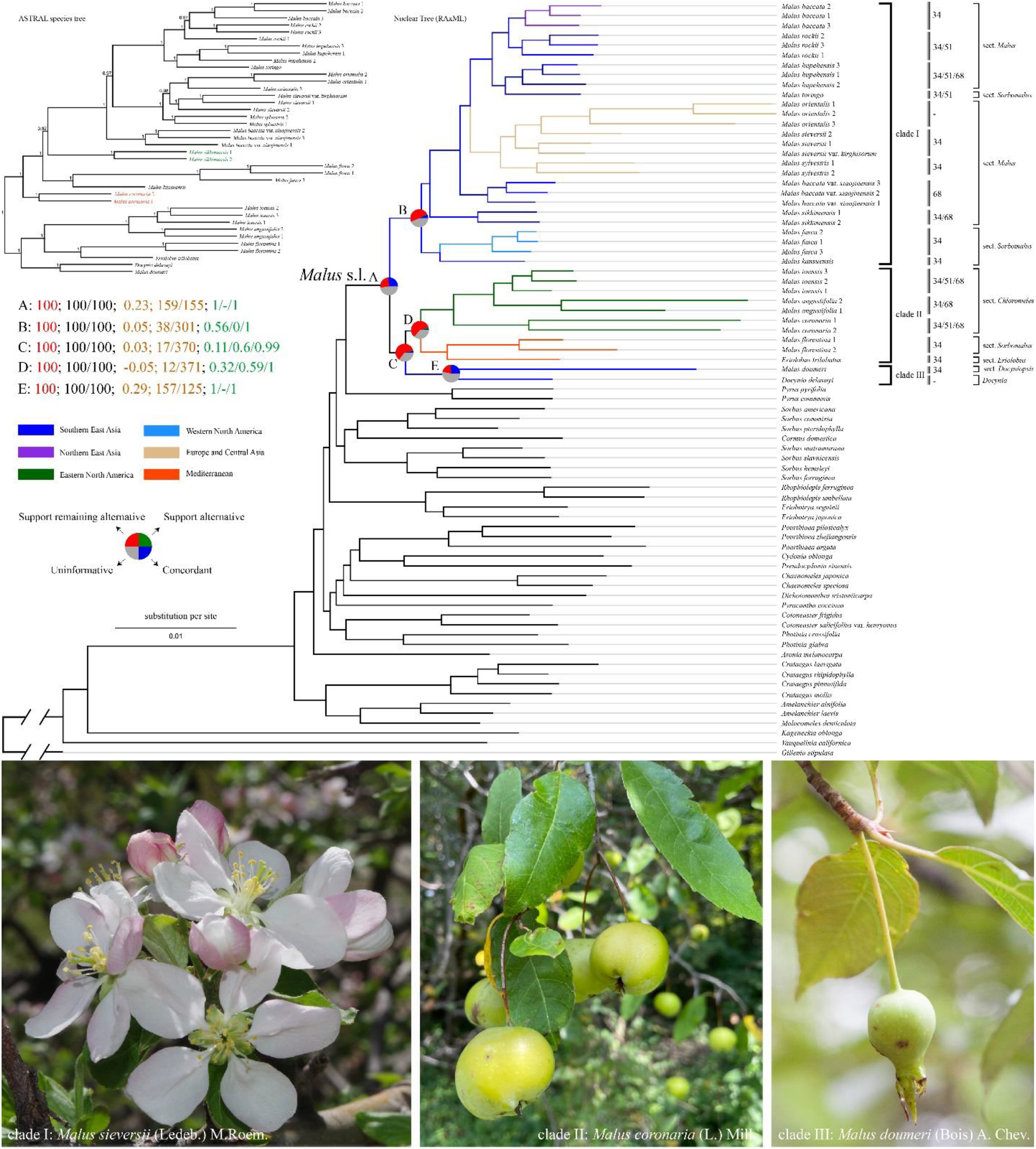
Maximum likelihood phylogeny of *Malus* s.l. in the framework of Maleae inferred from RAxML analysis of the concatenated 80%-sample dataset. Pie charts on the focused five nodes (A, B, C, D, and E) present the proportion of gene trees that support that clade (blue), the proportion that support the main alternative bifurcation (green), the proportion that support the remaining alternatives (red), and the proportion (conflict or support) that have < 50% bootstrap support (gray). All the other pie charts refer to Fig. S11 available from Dryad. The numbers (top left) indicate values associated with those nodes; they are bootstrap support values estimated from RAxML analysis (e.g., A: 100 labled by red; see Fig. S4 available on Dryad for all nodes BS), the SH-aLRT support and Ultrafast Bootstrap (UFBoot) support estimated from IQ-TREE2 (e.g., A: 100/100 labled by black; see Fig. S5 available on Dryad for all nodes support), the Internode Certainty All (ICA) score, the number of gene trees concordant/conflicting with that node in the species tree estimated from *phyparts* (e.g., 0.23; 159/155 labled by orange; see Fig. S11 available on Dryad for all values), and Quartet Concordance/Quartet Differential/Quartet Informativeness estimated from Quartet Samping analysis (e.g., 1/-/1 labled by green; see Fig. S10 available on Dryad for all scores). Branches are colored by their distribution, i.e., dark blue, Southern East Asia; purple, Northern East Asia; green, Eastern North America; light blue, Western North America; yellowish-brown, Europe and Central Asia; red, the Mediterranean. The sporophytic chromosome number is displayed to the right of each species label (see Table 2 for details). The Photo credits: clade I (*Malus sieversii*): Pan Li; clade II (*M. coronaria*): Richard G.J. Hodel; clade III (*M. doumeri*): Bin-Bin Liu.

Conflicted phylogenetic positions were detected in these nine phylogenetic trees (Fig. 3 and Supplementary Figs. S1-S9), in which the placements of *Malus coronaria* (L.) Mill. from eastern North America (Fig. 3: clade V) and the Asian *M. sikkimensis* (Wenz.) Koehne (Fig. 3: clade III) varied significantly among trees. All three species trees (Supplementary Figs. S3, S6, S9) estimated from ASTRAL-III supported the sister relationship between *M. coronaria* and a large clade consisting of all species in clade I, II, III, and IV (defined in Fig. 3) (i.e., *M. coronaria* bipartition A: Table 1), while the remaining six ML trees (Supplementary Figs. S1, S2, S4, S5, S7, S8) estimated from IQ-TREE2 and RAxML recovered *M. coronaria* as sister to the clade composed of another two eastern North American species (*M. angustifolia* (Aiton) Michx. and *M. ioensis* (Alph. Wood) Britton) (i.e., *M. coronaria* bipartition B: Table 1). Likewise, the phylogenetic position of *M. sikkimensis* showed incongruence among trees (Supplementary Figs. S1-S9). One bipartition supported by seven out of nine trees was the sister relationship between *M. sikkimensis* and the combined clade, including *M. baccata* var. *xiaojinensis* (M.H.Cheng & N.G.Jiang) Ponomar., *M. orientalis* Uglitzk., *M. sylvestris*, *M. sieversii*, *M. hupehensis* (Pamp.) Rehder, *M. toringo* (Siebold) de Vriese, *M. baccata*, and *M. rockii* Rehder (i.e., *M. sikkimensis* bipartition A: Table 1), while the other bipartition recovered by only two trees was the sister relationship between *M. sikkimensis* and the clade composed of two species (*M. fusca* (Raf.) C.K.Schneid. and *M. kansuensis* (Batalin) C.K.Schneid.) (*M. sikkimensis* bipartition B: Table 1). An edge-based phylogenomic support test implemented in *phyckle*^28^ on the 50%-sample dataset (600 genes all having *Gillenia stipulata* (Muhl. ex Willd.) Nutt. as outgroup) revealed that *M. coronaria* bipartition A was supported by more genes (445) than the *M. coronaria* bipartition B (155 genes). Meanwhile, the summed difference in log-likelihood scores supported the *M. coronaria* bipartition A (sum ΔlnL = 17028.36) over the bipartition B (sum ΔlnL = 4126.411). Although the extreme outlier genes were excluded (Table 1), both the number of genes (427:153 = 2.79 ≈ 3:1) and summed difference (14731.52:3876.897) supported the *M. coronaria* bipartition A. Similarly, two different bipartitions for the placements of *M. sikkimensis* were analyzed by *phyckle*. *Malus sikkimensis* bipartition B was supported by only 191 genes (sum ΔlnL = 3987.746) compared to the 409 genes of bipartition A (sum ΔlnL = 11675.01). Even with the outlier genes removed, *M. sikkimensis* bipartition B had more gene support (393, sum ΔlnL = 9168.462) than that of bipartition A (188, sum ΔlnL = 3647.953).

**Table 1.**
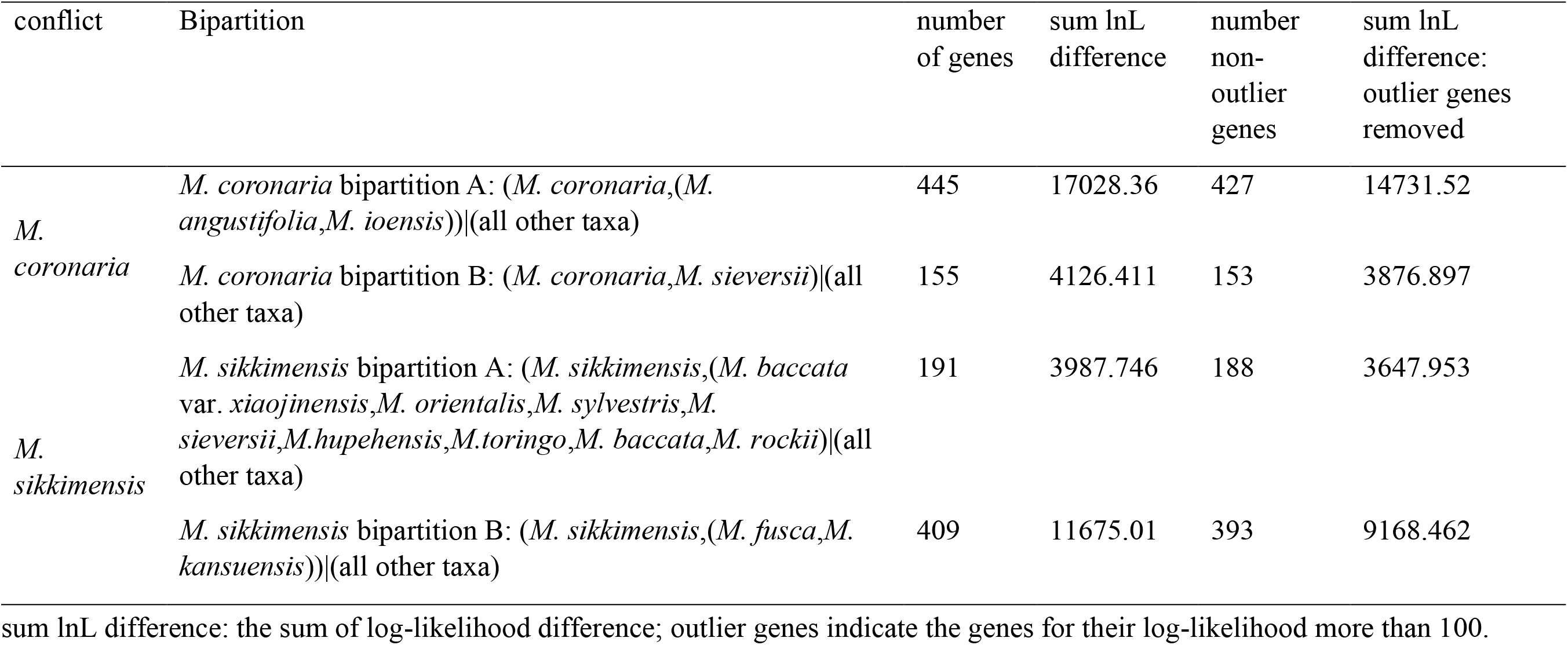
The result of *phyckle* analysis of detecting the gene dataset supporting each bipartition for *Malus coronaria* and *M. sikkimensis*. The result was estimated from the 50%-sample dataset.

| conflict | Bipartition | number<br>of genes | sum lnL<br>difference | number<br>non-<br>outlier<br>genes | sum lnL<br>difference:<br>outlier genes<br>removed |
| --- | --- | --- | --- | --- | --- |
| *M.*<br>*coronaria* | *M. coronaria* bipartition A: (*M. coronaria*,(*M. angustifolia*,*M. ioensis*))\|(all other taxa) | 445 | 17028.36 | 427 | 14731.52 |
|  | *M. coronaria* bipartition B: (*M. coronaria*,*M. sieversii*)\|(all other taxa) | 155 | 4126.411 | 153 | 3876.897 |
| *M.*<br>*sikkimensis* | *M. sikkimensis* bipartition A: (*M. sikkimensis*,(*M. baccata* var. *xiaojinensis*,*M. orientalis*,*M. sylvestris*,*M. sieversii*,*M. hupehensis*,*M. toringo*,*M. baccata*,*M. rockii*)\|(all other taxa) | 191 | 3987.746 | 188 | 3647.953 |
|  | *M. sikkimensis* bipartition B: (*M. sikkimensis*,(*M. fusca*,*M. kansuensis*))\|(all other taxa) | 409 | 11675.01 | 393 | 9168.462 |

sum lnL difference: the sum of log-likelihood difference; outlier genes indicate the genes for their log-likelihood more than 100.

The conflict analysis from *phyparts* showed that significant gene tree discordance was detected among nuclear genes regarding the placement of three major clades, and minimal informative gene trees supported each clade (Fig. 3 and Supplementary Fig. S11). In contrast, the QS result demonstrated that all these five nodes related to three major clades were confirmed with full support (1/-/1; i.e., all informative quartets support that lineage). Although Quartet Sampling (QS) confirmed the monophyly of *Malus* s.l. (node A) with full support, only 159 out of the 314 informative gene trees (50.6%) were concordant with this topology (ICA = 0.23). In contrast, only 56% of sampled informative quartets supported node B with only one alternative discordant topology (QS score = 0.56/0/1), but the *phyparts* result showed only 38 out of the 339 informative gene trees (11.2%; ICA = 0.05) supported this clade. Similarly, node E was supported by only 11% of informative quartets with a skewed frequency for alternative discordant topologies (QS score = 0.11/0.6/0.99), and the result of *phyparts* supported this clade with only 17 of the 387 informative gene trees (4.4%; ICA = 0.03). Node D was recovered in all nine trees (Supplementary Figs. S1-S9), while this was supported by only 12 of the 383 informative gene trees with counter-support from ICA (−0.05). Additionally, QS conflict analysis also showed the weak quartet support (0.32) with a skewed frequency for discordant topologies (0.59). In contrast, node E was recovered with full QS support (1/-/1) and 157 of the 282 informative gene trees (ICA = 0.29).

### Coalescence simulation and phylogenetic network analysis

The nuclear gene coalescent simulation analysis showed the distinguished empirical and simulated gene to gene distance distribution (Fig. 4h), suggesting that ILS alone can not explain most observed gene tree heterogeneity^61^. Of the 27-taxa dataset at the tribe level, the plot of pseudo-loglikelihood scores (Fig. 4g) showed that the optimal number of hybridization events inferred in the SNaQ network analysis was one (Fig. 4g: -ploglik = 5218.4), suggesting the hybrid origin of *Malus doumeri* between *Docynia delavayi* (γ = 0.794) and the ancestor of pome related members (i.e., the formerly Maloideae; γ = 0.206). As *h_max_* increased beyond one, the pseudo-loglikelihood increased slightly; therefore, we considered *h_max_* = 1 to be optimal for this 27-taxa dataset.

**Fig. 4.**
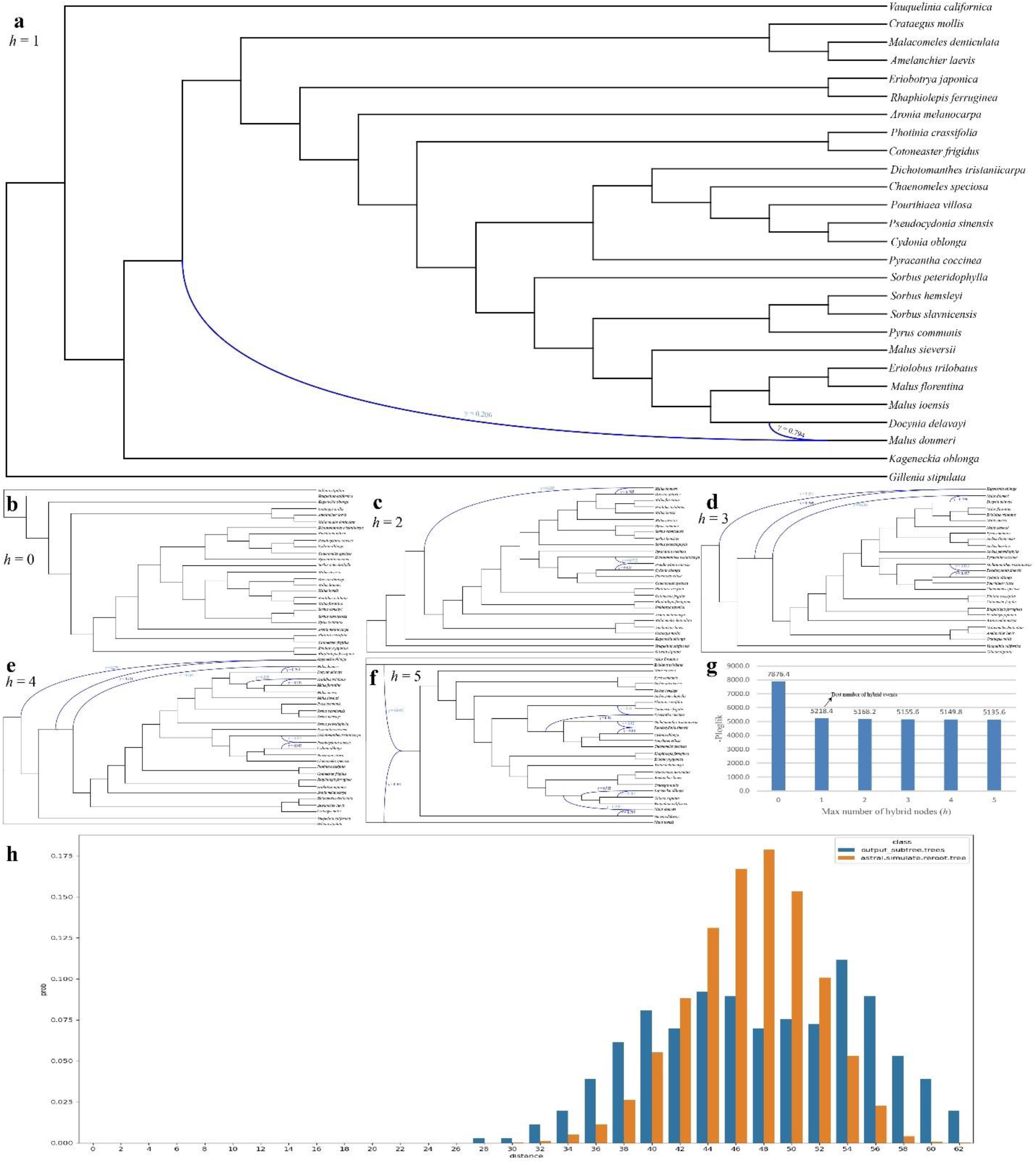
Coalescent simulation and phylogenetic network analysis from the 27-taxa sampling at the tribe level of Maleae. **a-f**, Species networks inferred from SNaQ network analysis with 1 to 5 maximum number of reticulations. **g**, The pseudo-loglikelihood scores (-ploglik) in a bar chart indicated that *hmax* = 1 was the optimal network (A). **h**, Distribution of tree-to-tree distances between empirical gene trees and the ASTRAL species tree, compared to those from the coalescent simulation. Blue curved branches indicate the possible hybridization event. Dark blue and light blue numbers indicate the major and minor inheritance probabilities of hybrid nodes.

The nuclear gene discordance analysis of the 14-taxa sampling dataset at the *Malus* level showed much overlap between the empirical and simulated distance distributions. This revealed that ILS alone might explain the observed phylogenomic discordance. Meanwhile, the optimal hybridization event inferred from SNaQ network analysis was also one (Fig. 5c: -ploglik = 491.5), because the score of pseudo-loglikelihood levels off when *h_max_* increased beyond one. *Malus coronaria* was 64.2% sister to a clade composed of *M. ioensis* and *M. angustifolia*, and 35.8% sister to one Central Asian species, *M. sieversii* (Fig. 5a).

**Fig. 5.**
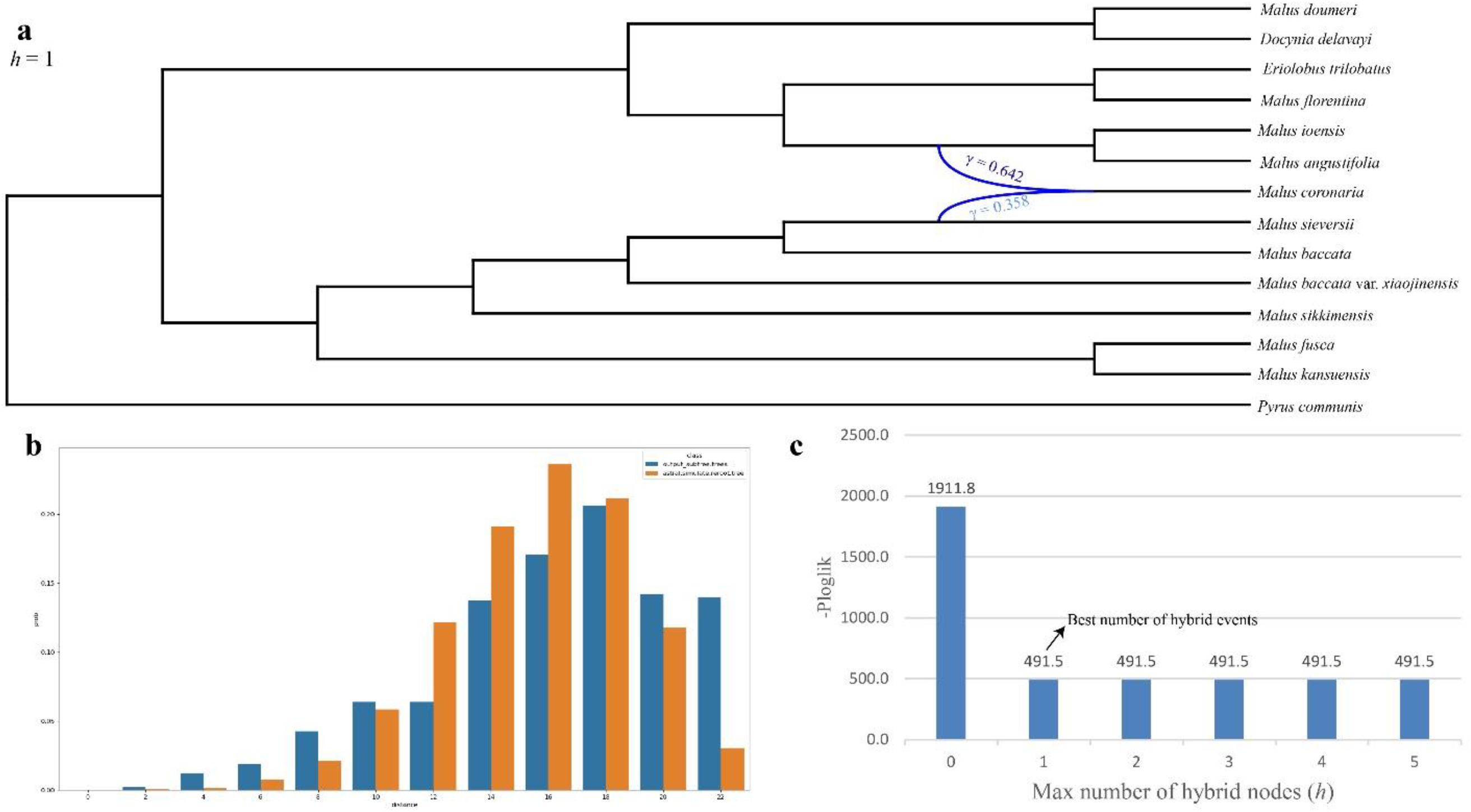
Coalescent simulation and phylogenetic network analysis from the 14-taxa sampling at the genus level of *Malus*. **a**, Species networks inferred from SNaQ network analysis with *hmax* = 1 as the optimal network. **b**, Distribution of tree-to-tree distances between empirical gene trees and the ASTRAL species tree, compared to those from the coalescent simulation. **c**, The pseudo-loglikelihood scores (-ploglik) of one to five maximum number of reticulations. Blue curved branches indicate the possible hybridization event. Dark blue and light blue numbers indicate the major and minor inheritance probabilities of hybrid nodes.

We used the filtered HyDe results (i.e., 0 < γ < 1) for detecting hybridization events. A total of 594 out of the 2448 hypotheses tested by HyDe showed significant evidence of a hybridization event (Supplementary Table S3), and nearly every species sampled in this study was involved in hybridization. The γ value for 350 of the 594 hypotheses was greater than 0.7 and less than 0.3, indicating ancient hybridization events, and only 244 γ values were close to 0.5 (0.3 < γ < 0.7), suggesting recent hybridization events. *Malus orientalis* has been involved in the most number of hybridization hypotheses (66), following with *M. coronaria* (54 ones) and *M. sikkimensis* (24 ones).

Due to the similar phylogenetic pattern between allopolyploidy and hybridization speciation, we summarized the chromosome count data for all species available from previous studies (Table 2, Fig. 3) for distinguishing the phylogenomic discordance from the two mechanisms. Generally, the chromosome distribution in *Malus* s.l. showed that allopolyploidy may have promoted the diversification of *Malus*. We did not find allopolyploidy in clade III. However, the various proportion of allopolyploidy cases was detected in clades I and II.

**Table 2.**
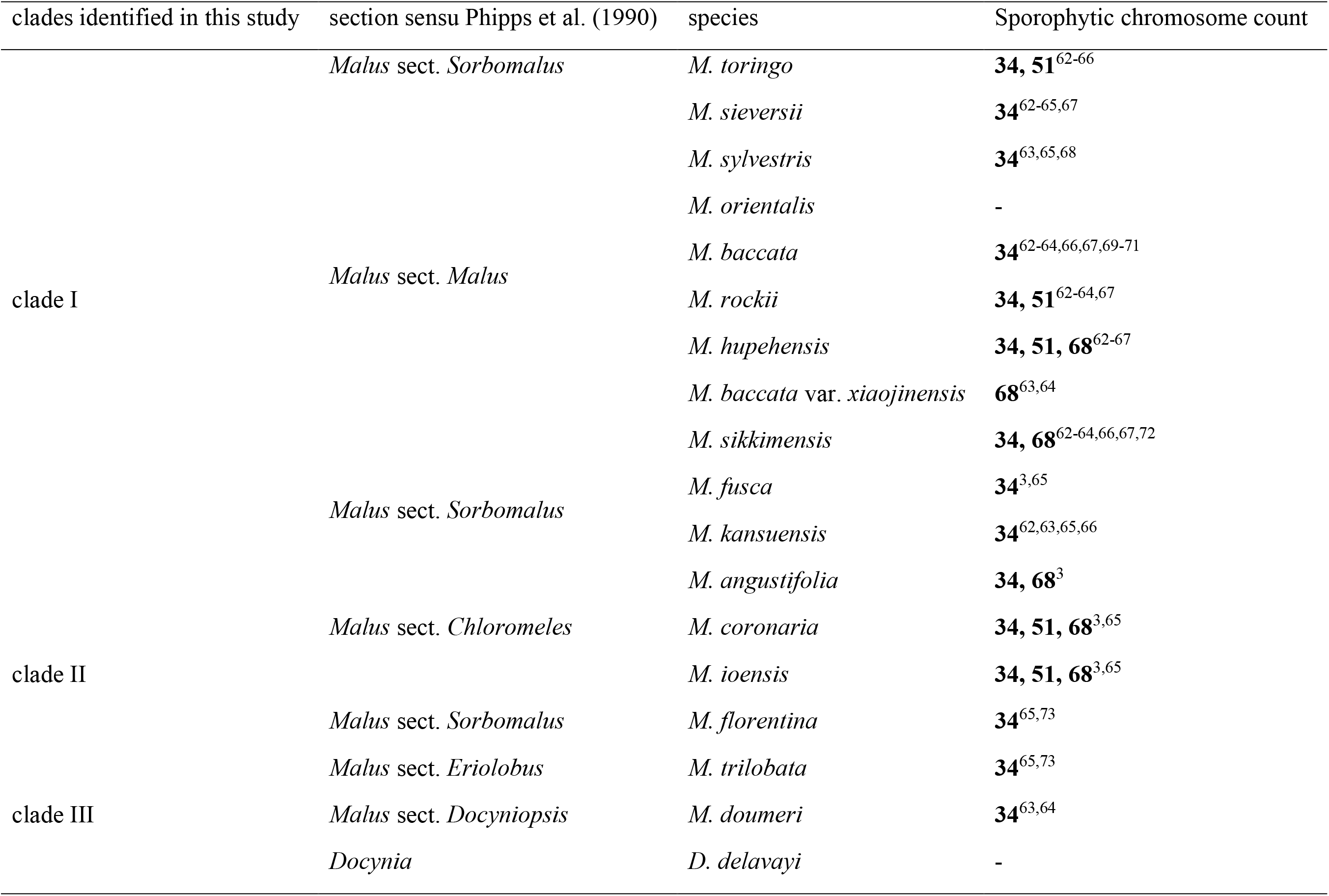
The sporophytic chromosome count number of *Malus* sampled in this study, along with their citations. These 18 species were grouped by Phipps et al. (1991)’s taxonomic system and the clades identified in this study.

### Plastid phylogenetic analysis

The final alignment from 80 plastid coding genes (CDSs) included 75 taxa and 80,799 bp in aligned length. All three phylogenetic trees from RAxML, IQ-TREE2, and ASTRAL-III recovered the same topology (Fig. 6 and Supplementary Figs. S12-S14). We presented the topology from RAxML herein and referred it to as the plastid phylogeny in the following context. Although the plastid result confirmed the three major clades in the nuclear phylogeny (Fig. 3), their relative phylogenetic position varied greatly (Fig. 6), and significant cytonuclear discordance showed between the plastid tree and the nuclear phylogeny (Fig. 7). The monophyly of *Malus* s.l. did not recover in the plastid phylogeny, and the eastern North American and Mediterranean species (clade II) were supported to be sister to *Pourthiaea* Decne. (Fig. 6). Due to the limited informative sites for each plastid coding gene, the *phyparts* resulted in nearly completely grey pies for each focal node, i.e., no or very few genes supported this node (Fig. 6 and Supplementary Fig. S16), suggesting the limited utility of *phyparts* in shallow phylogenies and/or plastid genes. By contrast, the QS conflict analysis showed full support for the five focal nodes (1/-/1).

**Fig. 6.**
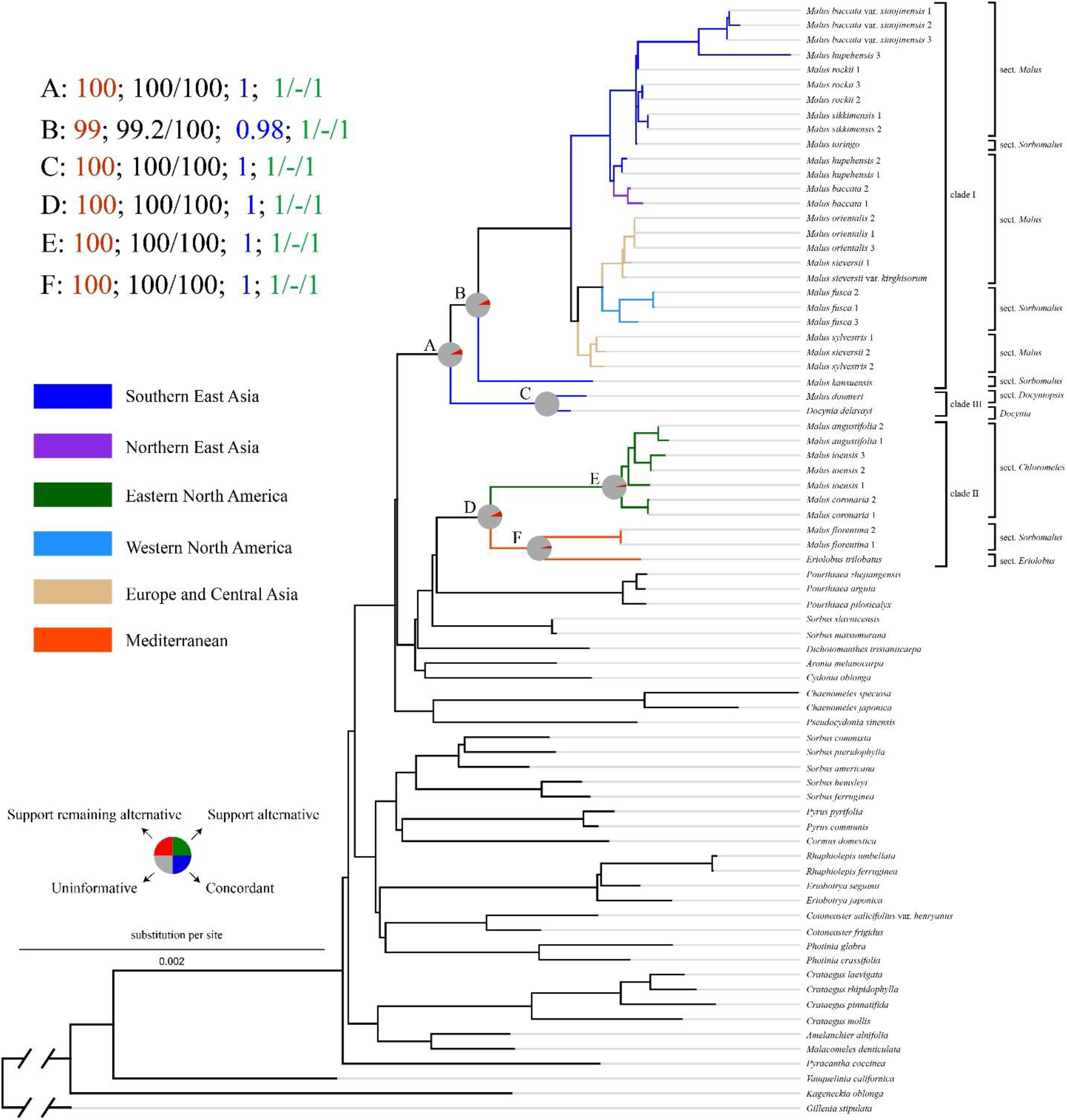
Maximum likelihood phylogeny of *Malus* s.l. in the framework of Maleae inferred from RAxML analysis of 80 plastid coding regions. Pie charts on the focused five nodes (A, B, C, D, E, and F) present the proportion of gene trees that support that clade (blue), the proportion that support the main alternative bifurcation (green), the proportion that support the remaining alternatives (red), and the proportion (conflict or support) that have < 50% bootstrap support (gray). All the other pie charts refer to Fig. S16, available from Dryad. The numbers (top left) indicate values associated with those nodes; they are bootstrap support values estimated from RAxML analysis (e.g., A: 100 labled by red; see Fig. S12 available on Dryad for all nodes BS), the SH-aLRT support and Ultrafast Bootstrap (UFBoot) support estimated from IQ-TREE2 (e.g., A: 100/100 labled by black; see Fig. S13 available on Dryad for all nodes support), the local posterior probability (LPP) estimated from ASTRAL-III (e.g., A: 1 labled by blue; see Fig. S14 available on Dryad for all LPP), and Quartet Concordance/Quartet Differential/Quartet Informativeness estimated from Quartet Samping analysis (e.g., 1/-/1 labled by green; see Fig. S15 available on Dryad for all scores). Branches are colored by their distribution, i.e., dark blue, Southern East Asia; purple, Northern East Asia; green, Eastern North America; light blue, Western North America; yellowish-brown, Europe and Central Asia; red, the Mediterranean.

**Fig. 7.**
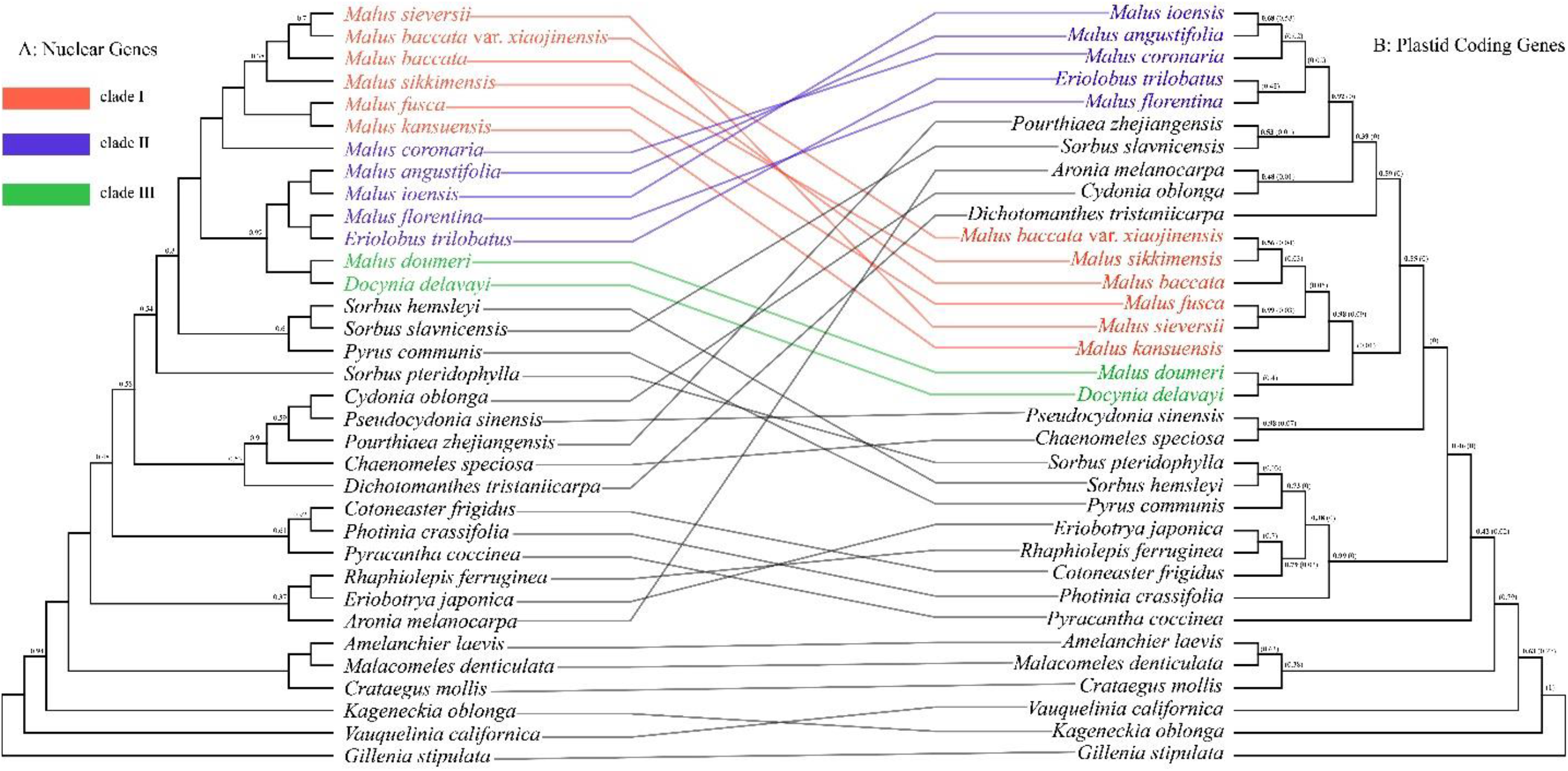
Tanglegram illustrating the cytonuclear discordance. Left: ASTRAL species tree based on SCN genes; right: plastid tree estimated from 80 coding genes. All nodes have maximum support (LPP = 1) unless rooted. Numbers in the brackets in the plastid tree show the contribution of ILS to the conflicts between nuclear and plastid gene trees based on the multispecies coalescent model.

### Dating and ancestral area reconstruction

The historical biogeographic analysis based on the SCN and plastid datasets using BEAST2 supported the East Asian origin of *Malus* s.l. The Northern Hemisphere disjunct distribution was through six (SCN data) or five (plastid data) dispersal events (Fig. 8). However, the phylogenetic dating analysis from SCN and the plastid dataset resulted in different age estimates. Generally, the overall age estimates in the nuclear data appeared to be older than those estimated from the plastid coding genes (Fig. 8 and Supplementary Figs. S17, S18). *Malus* s.l. originated from East Asia in the early Eocene, ca. 47.37 million years ago [Mya] in the SCN dating analysis, as compared to 42.16 Mya (95% highest posterior density (HPD) interval: 41.2-44.39 Mya: Supplementary Fig. S19) in the plastid analysis. The current disjunct distribution has been stabilized by the late Oligocene (26.42 Mya: 22.91-27.92 Mya: Fig. 8) based on the SCN result and by the early Miocene (15.14 Mya: 12.55-17.73 Mya: Supplementary Fig. S17) based on the plastid result. The eastern North American and Mediterranean clade (clade II: Fig. 3) was estimated to have originated from western North America in the middle Eocene [SCN: 43.58 Mya (41.2-44.39 Mya) and plastid coding genes: 41.2 Mya: 41.2-46.67 Mya], and then dispersed to the Mediterranean in the late Eocene [SCN: 39.61 Mya: 36.18-40.81 Mya; plastid coding genes: 35.77 Mya: 33.99-37.99 Mya) through North Atlantic Land Bridge (NALB). The SCN analysis showed that the eastern North American species originated from the extinct Western North American species in the late Eocene (34.97 Mya). In contrast, the plastid analysis resulted in the middle Miocene origin (14.18 Mya). Furthermore, our ancestral area reconstruction analysis from the SCN data showed that some *Malus* members dispersed back to western North America in the early Miocene (20.98 Mya: Fig. 8), which may be due to the climate oscillations then. However, our plastid result did not present this back dispersal (Fig. 8). Similarly, the Europe & Central Asia clade originated from an East Asian ancestor in the late Oligocene (SCN: 26.42 Mya: 22.91-27.92 Mya) and the middle Miocene (plastid coding genes: 15.14 Mya: 12.55-17.73 Mya). Contrastingly, the western North American lineage (*Malus fusca*) was estimated to have originated from East Asia in the early Oligocene (31.46 Mya: 26.08-34.92 Mya: Fig. 8) and migrated through the BLB by the SCN data, while it originated from Europe & Central Asia in the late Miocene (8.6 Mya: 6.46-10.91 Mya: Supplementary Fig. S17) and likely migrated through the NALB based on the plastid coding gene data.

**Fig. 8.**
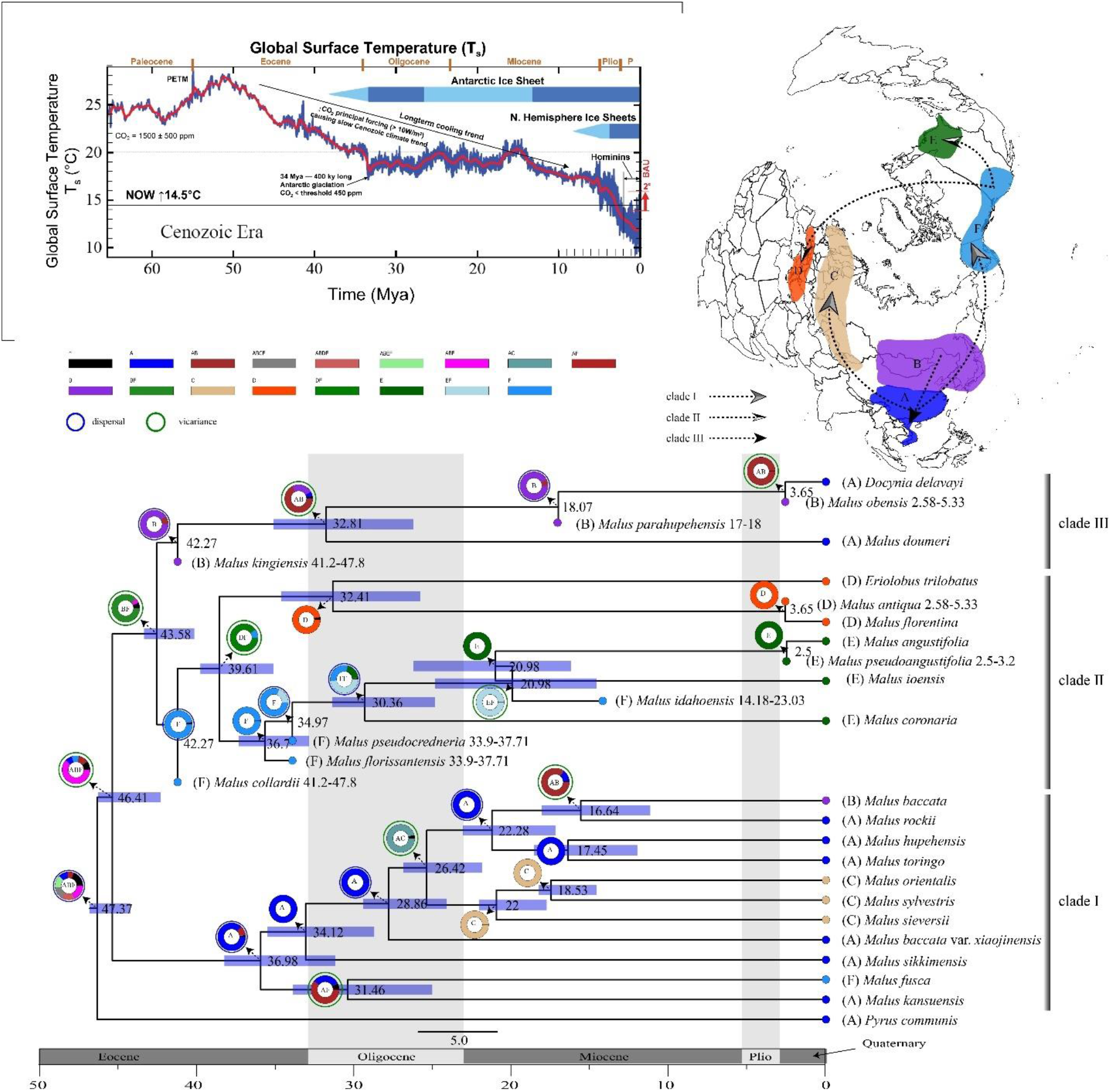
Dated chronogram for the apple genus *Malus* inferred from BEAST with the Fossilized Birth-Death process based on the 19-taxa nuclear dataset. Also shown is the ancestral area reconstruction using BioGeoBEARS implemented in RASP, with the colored key identifying extant and possible ancestral ranges (upper-right map), (A), Southern East Asia; (B), Northern East Asia; (C), Europe and Central Asia; (D), Mediterranean; (E), Eastern North America; (F), Western North America. The upper-left chart displays a global surface temperature, indicating the major global climate trends in the Cenozoic Era (adapted from https://www.alpineanalytics.com/Climate/DeepTime/WebDownloadImages/CenozoicTsGlobal-7.5w.600ppi.png).

## Discussion

This study integrated hundreds of SCN genes and plastomes to resolve the backbone of *Malus* s.l. The potential roles of ILS, hybridization, and allopolyploidy for the underlying phylogenetic discordance are herein evaluated. We also elucidate the biogeographic diversification patterns of the widespread disjunct distributions in the Northern Hemisphere. Our results support the paraphyly of the apple genus *Malus*, with *Docynia* nested within it, and recovered three strongly supported major clades within *Malus* s.l. except for the unstable phylogenetic placements of *M. coronaria* and *M. sikkimensis* in different trees. Furthermore, our coalescent simulation analysis demonstrated that ILS is not the sole or dominant cause of phylogenomic conflict, and other processes (e.g., hybridization and allopolyploidy) may have driven the discordance. The phylogenetic network analysis at the *Malus* and the Maleae levels supported the hybrid origin of *M. coronaria* and *M. doumeri*, and the further gene tree discordance analysis (e.g., *phyparts*, QS, *phyckle*, and HyDe) promoted the understanding of underlying conflict at some nodes. Below we will integrate several lines of evidence to discuss the potential causes of conflicts.

### The phylogenetic backbone and an early chloroplast capture event in *Malus* s.l

The taxonomic circumscription of *Malus* s.l. has been controversial due to the phylogenetic position of *Docynia*. With the highly distinctive numbers of ovules per locule (3-10 in *Docynia* vs. 2 in *Malus*), *Docynia* has been recognized as a separate genus by taxonomists historically^1, 5, 6, 10, 74–77^. The nuclear and plastid phylogeny from hundreds of SCN genes in our study supported the paraphyly of *Malus* s.s., with *Docynia* nested within it (Figs. 3, 4), and this was consistent with the phylogenetic inferences in several recent molecular studies^18, 21, 22^ (Fig. 1). *Docynia delavayi* (clade VIII: Fig. 3) was the sister of *Malus doumeri* (clade VII: Fig. 3), a species formerly treated in the *Malus* sect. *Docyniopsis* or the genus *Docyniopsis* (C.K.Schneid.) Koidz.. Several shared characteristics supported a close relationship between these species, i.e., cone-shaped non-adnate part of the ovaries, fully connate carpels, incurved and persistent calyx, numerous scattered sclereids throughout the flesh, juvenile leaves deeply lobed, and similar flavonoid chemistry^1, 5, 78^. Robertson et al. (1991) proposed to merge *Docynia* into *Docyniopsis* (= *Malus* sect. *Docyniopsis*) based on these shared characters. Given the more ovules per lucule (more than 2), *Docynia* was treated in *Cydonia* Mill. by Roemer (1847) and Wenzig (1883), and this treatment was not supported by our strongly supported nuclear and plastid phylogeny. Furthermore, several characters may easily distinguish *Docynia* from *Cydonia*, such as ovaries partially adnate to hypanthium in *Docynia* vs. fully adnate in *Cydonia*, styles fused at the base in *Docynia* vs. free in *Cydonia*, and stamens ca. 40 in *Docynia* vs. 25 in *Cydonia*^1, 5^. Hence, the above lines of evidence support a redefined circumscription of *Malus* s.l., by merging *Docynia* into *Malus*.

Our nuclear phylogeny recovered three major clades within *Malus* s.l.; however, cytonuclear discordance was detected for the placement of clade II, i.e., the combined eastern North American and Mediterranean species. Clade II was sister to clade III in the nuclear phylogeny (Fig. 3), while sister to another East Asian genus *Pourthiaea* in the plastid phylogeny (Figs. 4, 7). Several possible causes may explain the conflict between nuclear and plastid topologies, such as ILS, allopolyploidy, and hybridization^79^. Our coalescence simulation analysis showed that no simulated nuclear genes were concordant to the plastid tree, the well-supported incongruence of the clade II between the nuclear and plastid trees may not be explained by ILS (Fig. 7). Allopolyploidy could be excluded for explaining the discordance (Fig. 3, Table 2). Although the eastern Northern American species have been involved in diploidy, triploidy, and tetraploidy^3, 65^, the two Mediterranean species were consistently diploid^65, 73^. It is less likely that allopolyploidy has resulted in cytonuclear discordance.

Hybridization might be the underlying mechanism for explaining the conflicts between nuclear and plastid topologies (Fig. 7), especially in the context of the chloroplast capture hypothesis, which has been well illustrated by recent studies^21, 38, 80^. The chloroplast capture events have also been used to explain the topological conflicts in the AMP (*Amelanchier-Malacomelels-Peraphyllum*) clade of Maleae^21^. Furthermore, hybridization has played an essential role in the diversification of the apple tribe (Maleae)^5^. Our ancestral area reconstruction showed that the clade II originated from western North America in the middle Eocene (Fig. 8 and Supplementary Fig. S17). The shared plastomes between clade II and *Pourthiaea* implicated that the ancestor of clade II most likely captured the plastome of the ancestor of *Pourthiaea* in western North America, and the ancestor of *Pourthiaea* may have been widely distributed in the boreotropical flora of the Northern Hemisphere in Eocene^45, 81^. According to the chloroplast capture scenario of Liu et al. (2020a), we hypothesize that the ancestor of clade II might have been widely distributed in western North America, and served as the paternal parent of the hybrid. The pollen provider (the ancestor of the clade II) hybridized with the ancestor of *Pourthiaea* (the ovule provider), and formed the hybrid founder population. Subsequent backcrossings with the paternal parent promoted that the ancestor of clade II captured the complete plastome of the ancestor of *Pourthiaea*. Morphologically, clade II showed similarities to *Pourthiaea*, such as the densely scattered sclereids in the flesh of fruits^5, 82–84^.

### Ancient and recent events of hybridization and allopolyploidy drive the diversification of *Malus* s.l

Rapid diversification and reticulate evolution pose significant challenges for phylogenetic inference of *Malus* s.l. This study resolved three strongly supported clades using hundreds of SCN genes. However, underlying gene tree conflicts for most nodes showed the potential hybridization and allopolyploidy events in the early diversification of *Malus* s.l.^5, 85^.

Our phylogenetic network analysis at the Maleae level confirmed one hybridization event, i.e., the hybrid origin of *Malus doumeri* between *Docynia delavayi* and the ancestor of the pome-related members of Maleae (Fig. 4), supporting an ancient hybridization event in the evolutionary history of Maleae. *Malus doumeri* is a diploid^63, 64^ (Table1, 2*n* = 34); hence its origin did not involve an allopolyploidy event. Additionally, Our HyDe analysis detected 34 significant hybridization hypotheses that supported the hybrid origin of *M. doumeri*, and 25 out of 34 (γ < 0.3 and γ > 0.7) ones supported the ancient hybridization hypothesis.

Although the hybrid origin of *Malus coronaria* was estimated from the SNaQ analysis, the distribution of chromosome count data (2*n* = 34, 51, 68) indicated that allopolyploidy might have been involved in the speciation of *M. coronaria*, because the result of allopolyploidy event may have resembled that of hybridization^86^. In addition, our HyDe analysis showed that 27 of the 54 significant γ values supported the ancient hybridization events, and the same number of significant γ values (27) confirmed the recent hybridization events. However, an alternative hypothesis may explain this genetic admixture. With the wide naturalization of the European and the Central Asian species (*M. sylvestris*, *M. sieversii*, and *M. orientalis*) in eastern North America, the two individuals sampled in this study may represent a recent hybrid in the wild or horticulture. We need to test more samples of *M. coronaria* from its distributional range.

The phylogenetic conflicting positions of *Malus sikkimensis* among nine inferred topologies (Supplementary Figs. S1-S9) suggested its genetic heterogeneity, which also implicated by far more gene trees in conflict with the species tree than in concord (6/370) in the *phyparts* analysis and the limited QS quartets (0.4/0.76/0.99) (Fig. 3 and Supplementary Fig. S10). We suggest that allopolyploidy might have resulted in the phylogenetic discordance of *Malus sikkimensis* (Fig. 3) based on the uneven gene trees supporting each bipartition (Table 1). Other lines of evidence also support this hypothesis. The chromosome count of *M. sikkimensis* varied from diploid to tetraploid (Fig. 3, Table 2). Additionally, an equal number of significant ancient and recent hybridization events (12 : 12: Supplementary Table S2) estimated by HyDe analysis showed that frequent hybridization might have played an important role in the diversification of *M. sikkimensis*, which might have introgressed with other species in ancient and recent times.

Although the sister relationship between *Malus baccata* var. *xiaojinensis* and clade I (Fig 2) has been fully supported in all nine nuclear topologies (Supplementary Figs. S1-S9) and the nearly full support of the QS analysis (Supplementary Fig. S10), only 23 out of the 319 informative gene trees confirmed this relationship in the *phyparts* analysis (Supplementary Fig. S11). A series of previous studies have demonstrated exclusive apomixis for *M. baccata* var. *xiaojinensis*^87^. They may have derived from the hybridization events between *M. kansuensis* and *M. toringoides*^88^.

### An East Asian-western North American origin of *Malus* s.l. and its subsequent extinctions in the Eocene

Our historical biogeographic analysis inferred East Asian + western North America as the most likely ancestral area of *Malus* s.l. (Fig. 8). The common ancestor is postulated to have occupied a widespread East Asian-western North American range, consistent with the rich fossil records of clade II and III recovered in northeast Asia and western North America from Eocene to Pliocene (Table 3). However, we did not find any fossil records of clade I from southern East Asia, which may be due to the underexplored fossil discovery for this region. This result disagrees with the North American origin or the East Asian origin, as proposed by Nikiforova et al. (2013) and Jin (2014), respectively. The conflicted hypothesis on the *Malus* origin may be due to the absence of fossil records and the uneven sampling in the two prior analyses. Nikiforova et al. (2013)’s investigation did not include taxa of clade III (*Malus doumeri* and *Docynia delavayi*), and Jin (2014)’s study misidentified *Pseudocydonia sinensis* to be *Docynia delavayi*.

**Table 3.** Fossil records of *Malus*.

| Age | Fossil | fossil organ | geographic origin | reference |
| --- | --- | --- | --- | --- |
| Eocene (47.8-41.2 Mya) | *Malus collardii* Axelrod | leaves | North America (Thunder Mountain, Idaho, USA) | Axelrod, 1998 |
| Eocene (47.8-41.2 Mya) | *Malus kingiensis* Budants | leaves | Eurasia (Rebro Cape, western Kamchatka peninsula, Kamchatka Territory, Russian Federation) | Budantsev, 2006 |
| Eocene (37.71-33.9 Mya) | *Malus florissantensis* (Cockerell) MacGinitie | leaves | North America (Green River Formation, Florissant, Colorado, USA) | Cockerell, 1908; MacGinitie, 1953 |
| Eocene (37.71-33.9 Mya) | *Malus pseudocredneria* (Cockerell) MacGinitie | leaves | North America (Green River Formation, Florissant, Colorado, USA) | Cockerell, 1908; MacGinitie, 1969 |
| Miocene (23.03-14.18 Mya) | *Malus idahoensis* R.W.Br. | leaves | North America (G. W. Oliver coal mine, Salmon, Idaho, USA) | Brown, 1935 |
| Miocene (17 Mya)* | *Malus parahupehensis* J.Hsu & R.W.Chaney | leaves | NE China (Shanwang, Shandong, China) | Hsu & Chaney, 1940 |
| Pliocene (5.33-2.58 Mya) | *Malus obensis* M.G.Gorbunov | fruits | Eurasia (W Siberia) | Gorbunov, 1959 |
| Pliocene (5.33-2.58 Mya) | *Malus antiqua* Doweld = *Malus pulcherrima* Givulescu nom. illeg. | leaves | Europe (Chiuzbaia, Județul Maramureș, Romania) | Doweld (2018) |
| Pleistocene (3.2-2.5 Mya) | *Malus pseudoangustifolia* E.W.Berry | leaves | North America (right bank of Neuse River 4,5 miles above Seven Springs, Wayne County, North Carolina, USA) | Berry, 1926 |

\* The timing of the sedimentary sequence of the Shanwang Formation was estimated from $^{40}\text{Ar}/^{39}\text{Ar}$ analysis of the basalts<sup>102</sup>.

Our divergence time estimation suggested that three major clades of *Malus* diversified in the late Oligocene (43.58 Mya, 95% HPD interval: 41.2-44.39 Mya). Due to the decreased CO_2_ principal forcing and long-term cooling trend from the early Eocene, the high latitude *Malus* (clade III) from northern East Asia migrated to southern East Asia; the ancient western North American populations dispersed to eastern North America and the Mediterranean region (clade II: Fig. 8). Subsequent extinctions occurred in the northern cold area from the Eocene to the Quaternary because a series of fossil species from different eras have been discovered in northern East Asia (*Malus kingiensis* in the Eocene, *M. parahupehensis* in the Miocene, and *M. obensis* in the Pliocene: Table 3) and western North America (*M. collardii*, *M. florissantensis*, and *M. pseudocredneria* in the Eocene, and *M. idahoensis* in the Miocene: Table 3). These extinction events in northern East Asia may be related to the cooling events in geologic times, such as the Miocene cooling and drying occurred at approximately 15-10 Mya^89–91^ and the enhanced aridity at the middle latitudes of the Northern Hemisphere at about 8-7 Mya^92, 93^. The living species of clade III show preferences to cool habitats in the high altitudes of southern East Asia, suggesting the northern East Asian origin of clade III, such as *Malus doumeri* at 700-2400 m and *Docynia delavayi* at 1000-3000 m^2^. This distribution pattern has also been reported in many other angiosperm lineages, such as *Astilbe* Buch.-Ham. ex D.Don^94^, *Meehania* Britton ex Small & Vail^95^, *Mitchella* L.^96^, *Parthenocissus* Planch.^97^, and *Vitis* L.^98^. The extinctions of the early diverged *Malus* in western North America may be due to the increasing seasonality and drying spreading in the western Cordillera and cooling events into the Pleistocene^99–101^. The warm and moist environment in southern East Asia, eastern North America, and the Mediterranean promoted its survival in the refugia for *Malus* there, and the dispersal and vicariance events in the middle to late Eocene further facilitated its survival diversification across the Northern Hemisphere (Fig. 8).

## Materials & Methods

### Taxon sampling, library preparation, and deep genome skimming sequencing

Taxon sampling is designed to resolve the phylogenetic placements and relationships of major clades within *Malus* s.l. For the convenience of discussion, we followed the widely accepted taxonomic system proposed by Phipps et al. (1990), in which they recognized five sections. Due to the great economic importance of *Malus*, numerous hybrid cultivars have been used as crops and ornamentals, which have hybridized either within one section or between sections, and this made it difficult to infer the phylogenetic relationships between cultivated and wild species. Hence, we excluded the widely recognized artificial hybrid species in *Malus*, e.g., *M. × asiatica* Nakai, *M. × astracanica* hort. ex Dum.Cours., *M. × cerasifera* Spach, *M. × dawsoniana* Rehder, *M. × domestica* (= *M. pumila* Mill.), *M. × floribunda* Siebold ex Van Houtte, *M. × halliana*, *M. × micromalus*, *M. × heterophylla* Spach, *M. × magdeburgensis* Schoch ex Rehder, *M. × prunifolia* (Willd.) Borkh., *M. × sargentii* Rehder, *M. × scheideckeri* Späth ex Zabel, *M. × soulardii* (L.H.Bailey) Britton, *M. × spectabilis* (Aiton) Borkh., and *M. × sublobata* Rehder. We hence sampled 39 ingroup individuals representing 18 wild species (out of ca. 24^15^), representing all five sections and *Docynia* within *Malus* s.l. In addition, due to the potential biphyly of *Malus* based on complete plastomes^21, 103^, we sampled 38 outgroups across the tribes Maleae and Gillenieae to resolve the *Malus* phylogeny and identify possible events of cytonuclear discordance (Supplementary Table S1). We investigated 77 individuals in total, of which 27 species of deep genome skimming data were generated for this study (Supplementary Table S1).

Total genomic DNAs were extracted from silica-gel dried leaves or herbarium specimens using a modified CTAB (mCTAB) method^104^ in the lab of the Institute of Botany, Chinese Academy of Science (IBCAS) in China. The libraries were prepared in the lab of Novogene, Beijing, China using NEBNext^®^ Ultra^TM^ II DNA Library Prep Kit, and then paired-end reads of 2 × 150 bp were generated on the NovoSeq 6000 Sequencing System (Novogene, Beijing; 5-10 G data for each sample: Supplementary Table S1).

### Single-copy nuclear marker development

The SCN marker development followed the pipeline in Liu et al. (2021). Briefly, the coding regions of *Malus domestica* (GenBank assembly accession: GCA_000148765.2) were first input into MarkerMiner v.1.0^105^ to identify the putative single-copy genes. The resulting genes were then filtered by successively BLASTing^106–108^ against six available genomes [*Malus baccata* (accession: GCA_006547085.1), *M. domestica*, *Pyrus betulifolia* Bunge (accession: GCA_007844245.1), *P. bretschneideri* Rehder (accession: GCA_000315295.1), *P. ussuriensis* Maxim. × *P. communis* L. (accession: GCA_008932095.1), and *P. pyrifolia* (Burm.f.) Nakai (accession: GCA_016587475.1)] in Geneious Prime^109^, with the parameters settings in the Megablast program^110^ as a maximum of 60 hits, a maximum E-value of 1 × 10^-10^, a linear gap cost, a word size of 28, and scores of 1 for match and −2 for mismatch in alignments. We first excluded genes with mean coverage > 1.1 for alignments, which generally indicate potential paralogy of the genes and/or the presence of highly repeated elements in the sequences. The remaining alignments were further visually examined to exclude those genes receiving multiple hits with long overlapping but different sequences during the BLAST. It should be noted that the alignments with mean coverage between 1.0 and 1.1 were generally caused by the presence of tiny pieces of flanking intron sequences in the alignments. These fragments were still accepted as a SCN gene here. After filtering, the remaining genes were used as references in the following gene assembly. The baits *in silico* could be available from the Dryad Digital Repository (Data 1): https://doi.org/10.5061/dryad.2jm63xsq5.

### Data processing and the assembly of single-copy nuclear genes

Read processing and assembly followed the pipeline in Liu et al. (2021). Generally, we used Trimmomatic v. 0.39^111^ for quality trimming and adapter clipping with the parameters (ILLUMINACLIP:TruSeq3-PE.fa:2:30:10:1:true LEADING:3 TRAILING:3 SLIDINGWINDOW:4:15 MINLEN:36). Then, the results were quality-checked with FastQC v. 0.11.9^112^. The number of clean reads after trimming was also calculated here for comparison (Supplementary Table S1). We used the HybPiper pipeline v. 1.3.1^113^ for targeting SCN genes with default settings; BWA v. 0.7.1^114^ to align and distribute reads to target genes; SPAdes v. 3.15.0^115^ with a coverage cutoff value of 8 to assemble reads to contigs; and Exonerate v. 2.2.0^116^ to align assembled contigs to target sequences and determine exon-intron boundaries. Python and R scripts included in the HybPiper pipeline^113^ were used to retrieve the recovered gene sequences, summarize and visualize the recovery efficiency.

### Nuclear datasets construction and phylogenetic analysis

Sequences for each SCN were aligned in MAFFT v. 7.480^117^ with options “--localpair --maxiterate 1000”. Due to the variable sequencing coverage of each sample in this study, we employed three steps to remove the poorly aligned regions. We used trimAL v. 1.2^118^ to trim the alignment of each SCN, in which all columns with gaps in more than 20% of the sequences or with a similarity score lower than 0.001 were removed. Given the low-quality assembly in some sequences, Spruceup^119^ was used to discover, visualize, and remove outlier sequences in the concatenated multiple sequence alignments with the window size 50 and overlap 25. Because the Spruceup algorithm works better, the more data it has, we concatenated all the SCN gene alignments with AMAS v. 1.0^120^ before running Spruceup. We also used AMAS v. 1.0^120^ to split the processed/trimmed alignment back into single-locus alignments. The resulting alignments for each SCN were trimmed again using trimAL v. 1.2^118^ with the same parameters described above. Thirdly, we excluded the sequences less than 250 bp in each alignment with our customized python script (exclude_short_sequences.py, which can be available from Dryad Digital Repository https://doi.org/10.5061/dryad.2jm63xsq5) for decreasing the effect of missing data, because the short sequences in each alignment have few informative sites for the following coalescent-based species tree inference. The resulting SCN genes were used to infer individual ML gene trees using RAxML 8.2.12^121^ with a GTRGAMMA model and the option “-f a” and 200 BS replicates to assess clade support for each SCN. TreeShrink v. 1.3.9^122^ was used for detecting abnormally long branches in each tree with the default false positive error rate α = 0.05 and per-species mode. The shrunk trees and sequences have been used for the following phylogenetic inference, and hereafter these resulted sequences were referred to as “clean genes”.

We generated three different datasets to reconstruct the phylogeny to account for the effect of missing data in each SCN gene: (1) 50%-sample dataset: each SCN gene with at least 900 bp and more than 50% samples (≥ 39 individuals); (2) 80%-sample dataset: each SCN gene with at least 900 bp and more than 80% samples (≥ 62 individuals); (3) all-sample dataset: each SCN with at least 900 bp and more than 100% samples (77 individuals). These three datasets can be available from the Dryad Digital Repository (Data 2, 3, 4): https://doi.org/10.5061/dryad.2jm63xsq5. We used both concatenated and coalescent-based methods for phylogenetic inference of each dataset. We used PartitionFinder2^123, 124^ to estimate the best-fit partitioning schemes and/or nucleotide substitution models under the corrected Akaike information criterion (AICc) and linked branch lengths, as well as with rcluster^125^ algorithm options for the nuclear dataset. The resulting partitioning schemes and evolutionary models were used for the following Maximum Likelihood (ML) tree using IQ-TREE2 v. 2.1.3^126^ with 1000 SH-aLRT and the ultrafast bootstrap replicates and RAxML 8.2.12^121^ with GTRGAMMA model for each partition and clade support assessed with 200 rapid bootstrap (BS) replicates. The shrunk trees from TreeShrink^122^ were used to estimate a coalescent-based species tree with ASTRAL-III (Zhang et al., 2018) using local posterior probabilities (LPP)^127^ to assess clade support. Each of the gene trees was rooted, and low support branches (≤ 10) were collapsed using Newick Utilities^128^ or *phyx*^129^ since collapsing gene tree nodes with BS support below a threshold value will help to improve accuracy^130^. In total, nine phylogenies were generated for topological comparison, and these nine trees are available from the Dryad Digital Repository: https://doi.org/10.5061/dryad.2jm63xsq5.

### Detecting and visualizing nuclear gene tree discordance

To explore the discordance among gene trees, we employed *phyparts* v. 0.0.1^29^ to calculate the conflicting/concordant bipartitions by comparing the nuclear gene trees against the ML tree inferred from RAxML with a BS threshold of 50 (i.e., gene-tree branches/nodes with less than 50% BS were considered uninformative) for filtering out poorly supported branches, thus alleviating noise in the results of the conflict analysis^29^. We also used the internode certainty all (ICA) value that resulted from *phyparts* to quantify the degree of conflict on each node of a species tree given individual gene trees^131^. *Phyparts* results were visualized with *phypartspiecharts.py* (by Matt Johnson, available from https://github.com/mossmatters/MJPythonNotebooks/blob/master/phypartspiecharts.py). Furthermore, in order to distinguish lack of support from conflicting support in the species tree, we conducted Quartet Sampling (QS)^41^ analysis with 100 replicates and the log-likelihood cutoff 2. The QS method subsamples quartets from the input tree and alignment to assess the confidence, consistency, and informativeness of internal tree relationships, and the reliability of each terminal branch, and then four values are given in this analysis: QC = Quartet Concordance, QD = Quartet Differential, QI = Quartet Informativeness, and QF = Quartet Fidelity. The QS result was visualized with plot_QC_ggtree.R (by ShuiyinLIU, available from https://github.com/ShuiyinLIU/QS_visualization). Both the *phyparts* and QS results can provide alternative evidence for evaluating the discordance between gene trees.

Comparing the nine topologies inferred above, we found two species (*Malus coronaria* and *M. kansuensis*) with conflicting phylogenetic positions among trees (referring to the result below). We used the “alternative relationship test” in *phyckle*^28^ to investigate conflicting bipartitions and discover the gene trees supporting each bipartition. Considering the complex origin of *Malus coronaria*, we employed the two bipartitions supported by Phylonetworks analysis (see Fig. 4 & Table 1 in Results). Two or more user-specified alternative bipartitions could be used as a constraint to infer gene trees. Arbitrarily, we set the cutoff of ΔlnL > 100 as the outlier genes^132^. The resulting gene dataset supporting each bipartition was then used to estimate phylogenetic inference based on the concatenated (IQ-TREE2 and RAxML) and coalescent-based methods (ASTRAL-III) mentioned above. The resulting 12 trees could be available from the Dryad Digital Repository: https://doi.org/10.5061/dryad.2jm63xsq5.

### Coalescence simulation and phylogenetic network estimation

To measure the goodness-of-fit of the coalescent model with ILS explaining the gene tree discordance adequately, we performed a coalescent simulation analysis following the methods in previous studies^34, 133–135^. Briefly, if the simulated gene trees based on the coalescent model correspond well to the empirical gene trees, the gene tree discordance may be explained by ILS. Given the calculation of gene tree distances, we subsampled 27 species representing each major clade in the phylogenetic framework of the tribe Maleae. The function “sim.coaltree.sp” implemented in the R package Phybase v. 1.5^136^ has been used to simulate 10,000 gene trees with the multispecies coalescent (MSC) model.

Hybrid Detector (HyDe) can be used to detect hybridization using phylogenetic invariants arising under the coalescent model with hybridization^137^. We sampled 40 taxa, including all 39 *Malus* s.l. individuals and one outgroup (*Pyrus communis*) to detect the possible hybridization events within *Malus* s.l. This dataset can be available from the Dryad Digital Repository (Data 6): https://doi.org/10.5061/dryad.2jm63xsq5. The γ denotes the inheritance probability of parent 1 (P1), while 1 - γ would be the probability of the hybrid population being sister to parent 2 (P2). Generally, significant γ values close to 0.5 indicate a recent hybridization event; significant γ values closer to 0 or 1 indicate an ancient hybridization event remained in the extant species. We herein set the γ threshold at 0.3 and 0.7, which followed convention^132^.

To explore the possibility of reticulation as a cause of discordance in the apples and their allies, we employed the Species Networks applying Quartets (SNaQ) method^43^ implemented in the software PhyloNetworks 0.14.0^44^, which explicitly accommodates introgression/gene flow and ILS. Given the computational limitation of PhyloNetworks, we used two datasets to test for hybridization events, i.e., 27-taxa sampling at the tribe level of Maleae and 14-taxa sampling at the genus level of *Malus*. These two datasets are available from the Dryad Digital Repository (Data 7 & 8): https://doi.org/10.5061/dryad.2jm63xsq5. Considering that *Malus* member*s* have been reported to have hybridized with many genera in Maleae, e.g., *Aria* (Pers.) Host and *Torminaria* M.Roem.^5^, we sampled 27 individuals, including five species representing each major clade in *Malus* s.l. and 22 outgroup species in Maleae. This taxon sampling scheme represents a reasonable compromise between taxonomic coverage and computational cost. The 27-taxa dataset construction followed the method described above (i.e., Nuclear dataset construction and phylogenetic analysis). The best trees generated from RAxML were used to estimate the quartet concordance factors (CFs), representing the proportion of genes supporting each possible relationship between each set of four species. The resulting CFs and the ASTRAL species tree were used as initial input to run SNaQ analysis (*h* = 0), and the resulting best network was used as starting topology to run the next *h* value (*h* + 1), and so on. We investigated *h* values ranging from 0 to 5 with 50 runs in each *h* for estimating the best phylogenetic network. Each run generated a pseudo-deviance score: a value for fitting the network to the data, and estimated the inheritance probabilities (i.e. the proportion of genes contributed by each parental population to a hybrid taxon) for each network. Similarly, we also sampled 14 species, including 13 *Malus* species and one outgroup (*Pyrus communis*), to test the hybridization events among *Malus* members. The method followed that of the 27-taxa sampling dataset mentioned above. The best network was visualized using Dendroscope v 3.7.4^138^.

### Plastome assembly, annotation, phylogenetic analysis, and cytonuclear discordance

A two-step strategy was used for obtaining high-quality chloroplast genomes. NOVOPlasty v. 4.3.1^139^ was applied first to assemble the plastomes with high-quality raw data. Then we used the successive assembly approach^140^, combining the reference-based and the *de novo* assembly methods to assemble the remaining low-quality samples. With the *de novo* assembly and a seed-and-extend algorithm, NOVOPlasty was the least laborious approach and resulted in accurate plastomes; however, this program needs sufficient high-quality raw reads without gaps to cover the whole plastome. The whole plastomes assembled from NOVOPlasty then could be used as references for assembling the remaining samples. The successive method provided an excellent approach to obtaining relatively accurate and nearly complete plastomes with or without gaps from lower-coverage raw data. Due to the sensitivity of Bowtie2 v. 2.4.2^141^ to the reference, this successive method needs a closely related reference sequence with increased time and RAM requirements. Several recent studies have described the procedure in detail^21, 60, 103, 140, 142, 143^. All assembled plastomes have been submitted to GenBank with the accession numbers listed in Supplementary Table S1.

The assembled plastid genomes from the low-coverage and high-coverage datasets were annotated using PGA^144^ with a closely related plastome (MN062004: *Malus ioensis*) downloaded from GenBank as the reference, and the results of automated annotation were checked manually. The coding sequences of plastomes were translated into proteins to manually check the start and stop codons in Geneious Prime^109^. The custom annotations in the GenBank format were converted into the FASTA format, and five-column feature tables file required by NCBI submission using GB2sequin^145^.

Given the considerable variation among plastid introns at the level of Maleae, we extracted 80 coding genes (CDSs) using Geneious Prime^109^, and these CDSs were aligned by MAFFT v. 7.475^117^ with default parameters, respectively. This dataset with 77 samples and 80 plastid coding genes is available from the Dryad Digital Repository (Data 5): https://doi.org/10.5061/dryad.2jm63xsq5. The best-fit partitioning schemes and/or nucleotide substitution models for each dataset were estimated using PartitionFinder2^123, 124^, under the corrected Akaike information criterion (AICc) and linked branch lengths, as well as with rcluster^125^ algorithm options. The partitioning schemes and evolutionary model for each subset were used for the downstream phylogenetic analysis. Like the nuclear analysis, we estimated the ML tree by IQ-TREE2 v. 2.1.3^126^ with 1000 SH-aLRT and the ultrafast bootstrap replicates and RAxML 8.2.12^121^ with GTRGAMMA model for each partition and clade support assessed with 200 rapid BS replicates. We also used ASTRAL-III^130^ for estimating a coalescent-based species tree. These three trees are available from the Dryad Digital Repository: https://doi.org/10.5061/dryad.2jm63xsq5. Cytonuclear discordance at various levels was detected in *Malus* s.l. (Fig. 7); coalescent simulation was also employed for evaluating the importance of ILS in explaining cytonuclear discordance^35, 135^. We used the R package Phybase v. 1.5 to simulate 10,000 gene trees based on the ASTRAL species tree from SCN genes under MSC model. We used *Phyparts* v. 0.01^29^ to explore conflicts among the simulated gene trees and plastid tree, and the proportion of gene tree concordance compared to the plastid genome tree was used to qualify the discordance.

### Dating and ancestral area reconstruction

We aim to estimate the age of divergence of the three major clades identified from hundreds of SCN genes and 80 plastid coding genes in *Malus* s.l. The fossils used in this study are listed in Table 3. *Malus obensis* and the clade *Malus doumeri* - *Docynia delavayi* were grouped together because of the fruit similarity. *Malus parahupehensis* was thought to be similar to the living species *M. hupehensis*; however, this fossil species showed more similarities to *M. doumeri* because of the dentate margin and parallel, craspedodromous venation^15^. Thus, we grouped *M. parahupehensis*, *M. doumeri, M. obensis*, and *Docynia delavayi* as monophyletic. The earliest fossil, *Malus kingiensis,* was found in the middle Eocene from the western Kamchatka peninsula and can not be assigned to any living lineages of *Malus* s.l. based on the morphology; thus, this fossil species likely represented the stem clade of *Malus doumeri* and *Docynia delavayi*. However, only one fossil, *Malus antiqua*, was found in Europe, and this fossil with deeply lobed leaves was similar to the Mediterranean species. Therefore, we grouped this fossil with the two Mediterranean living species (*M. florentina* and *Eriolobus trilobatus*). Five fossil species were described from North America, especially the abundant fossil records in western North America. More or less deeply lobed leaves characterized all these leaf fossils, significantly distinct from the living western North American species (*Malus fusca*). They showed more similarities to the eastern North American and Mediterranean species, and these species were thus grouped together. Additionally, leaf fossil of *Malus* or *Pyrus* from the Republic site, Washington^146^ was used to constrain the divergence between *Malus* and *Pyrus* at 46-44 Mya.

We ran the dating analyses based on the SCN and plastid datasets to test the divergence time differences with significant cytonuclear discordance. Given the intensive computational burden of dating analysis using BEAST2, we employed a 19-taxa dataset with only one individual for each species in *Malus* s.l. and *Pyrus communis* as the outgroup, and this dataset is available from the Dryad Digital Repository (Data 13): https://doi.org/10.5061/dryad.2jm63xsq5. The divergence time estimation was run under a GTR model with a gamma rate inferred from PartitionFinder2^123, 124^, an uncorrelated lognormal relaxed clock^147^, and the fossilized birth-death model^148, 149^. Markow Chain Monte Carlo (MCMC) chains were run for 100,000,000, sampling every 20,000 generations in five parallel jobs. We used the LogCombiner v1.10 to combine log and tree files from the five independent runs of BEAST. The MCMC trace file was analyzed in Tracer v1.7.1^150^. Maximum credibility trees were generated in TreeAnnotator v1.10, and FigTree v1.4.4 visualized the MCC tree.

To test the ancestral areas of three major clades of *Malus*, we conducted the ancestral area construction using BioGeoBEARS v. 1.1.1^151^ implemented in RASP v. 4.2^152^. Geological evidence suggests that an aridity barrier existed from the western-most part of China to the eastern Asian coast from the Paleogene to the Miocene. It has been hypothesized to have acted as a climate barrier between these two regions^153^; thus, we subdivided East Asia into the northern and the southern areas^154–159^. Six biogeographic areas were defined across the distribution of *Malus* s.l.: (A), Southern East Asia; (B), Northern East Asia; (C), Europe and Central Asia; (D), Mediterranean; (E), eastern North America; (F), western North America. The MCC tree summarized by TreeAnnotator was used as input of RASP. The *maxarea* was set to six, i.e., the number of potential areas of a hypothetical ancestor was restricted to a maximum of six regions. The model with the highest AICc_wt value has been chosen as the best model.

## Conclusions

We resolved the phylogenetic backbone of *Malus* s.l. using 785 nuclear loci (77 taxa) and 80 plastid coding genes (75 taxa). The nuclear phylogeny supported the monophyly of *Malus* s.l. (including *Docynia*) and three strongly supported major clades within the genus. However, widespread gene tree conflicts among nuclear gene trees indicated the complicated evolutionary history of *Malus*, and ILS, hybridization, and allopolyploidy have played an important role in the evolution of *Malus*, explaining this cytonuclear discordance. We detected a deep hybridization event involving *Malus doumeri* as a hybrid between the ancestor of pome-beared species and *Docynia delavayi*, and a recent hybridization event (*M. coronaria*) between *M. sieversii* and a taxon of the clade of *M. ioensis* and *M. angustifolia*. However, our plastid result recovered the biphyly of *Malus* s.l., with the combined eastern North American and the Mediterranean clade sister to the East Asian genus *Pourthiaea*. The well-supported cytonuclear discordance could be best explained by the chloroplast capture event that occurred in western North America in the Eocene. The phylogenomic case study of the apple genus implicated that multiple methods accounting for ILS and gene flow can help untangle complex phylogenetic relationships among species, and concatenation method or methods only accounting for ILS (coalescent-based method) are biased and not appropriate for phylogenetic inferences in lineages with a highly complex evolutionary history. Our historical biogeographic analysis without fossil species in northern East Asia and western North America resulted in the East Asian origin, contrastingly that involving fossil and living species supported a widespread East Asian-western North American origin of *Malus* s.l., followed by subsequent extinction events in northern East Asia and western North America in the Eocene, and this indicated that integrating fossil and living species could promote the more accurate estimation of dating analysis, as well as ancestral area reconstruction^160^. The robust phylogenomic framework for the apple genus should provide an evolutionary basis for the breeding and crop improvement of apples and their close relatives. The study also represents an excellent case for utilizing the recently proposed deep genome skimming approach for obtaining nuclear and organelle genes for robust phylogenetic reconstructions.

## Acknowledgments

All computational analyses were conducted on the Smithsonian Institution High Performance Computing Cluster (SI/HPC, “Hydra”: https://doi.org/10.25572/SIHPC). National Natural Science Foundation of China supports this research (Grant No.: 32000163 & 31620103902).

## Author contributions

B.B.L. designed and led the project. D.Y.H. and J.W. supervised the study. B.B.L. carried out the phylogenomic analyses and wrote the draft manuscript. C.X. performed the experiment of deep genome skimming. C.R., M.K., R.H., J.H., and W.B.Z. participated in the phylogenetic and biogeographic analyses. G.Z.Q. provided suggestions on structuring the paper. C.H.H. and H.M. provided part of the transcriptomic data. All the authors contributed to the writing and interpretation of the results, and approved the final manuscript.

## Data availability

Raw reads have been deposited in the NCBI Sequence Read Archive (BioProject PRJNA759205). The customized scripts for SCN gene analysis, tree files, pre-and post-filtered alignments for all analyses are available from the Dryad Digital Repository: https://doi.org/10.5061/dryad.2jm63xsq5.

## Supplementary files

**Fig. S1.**
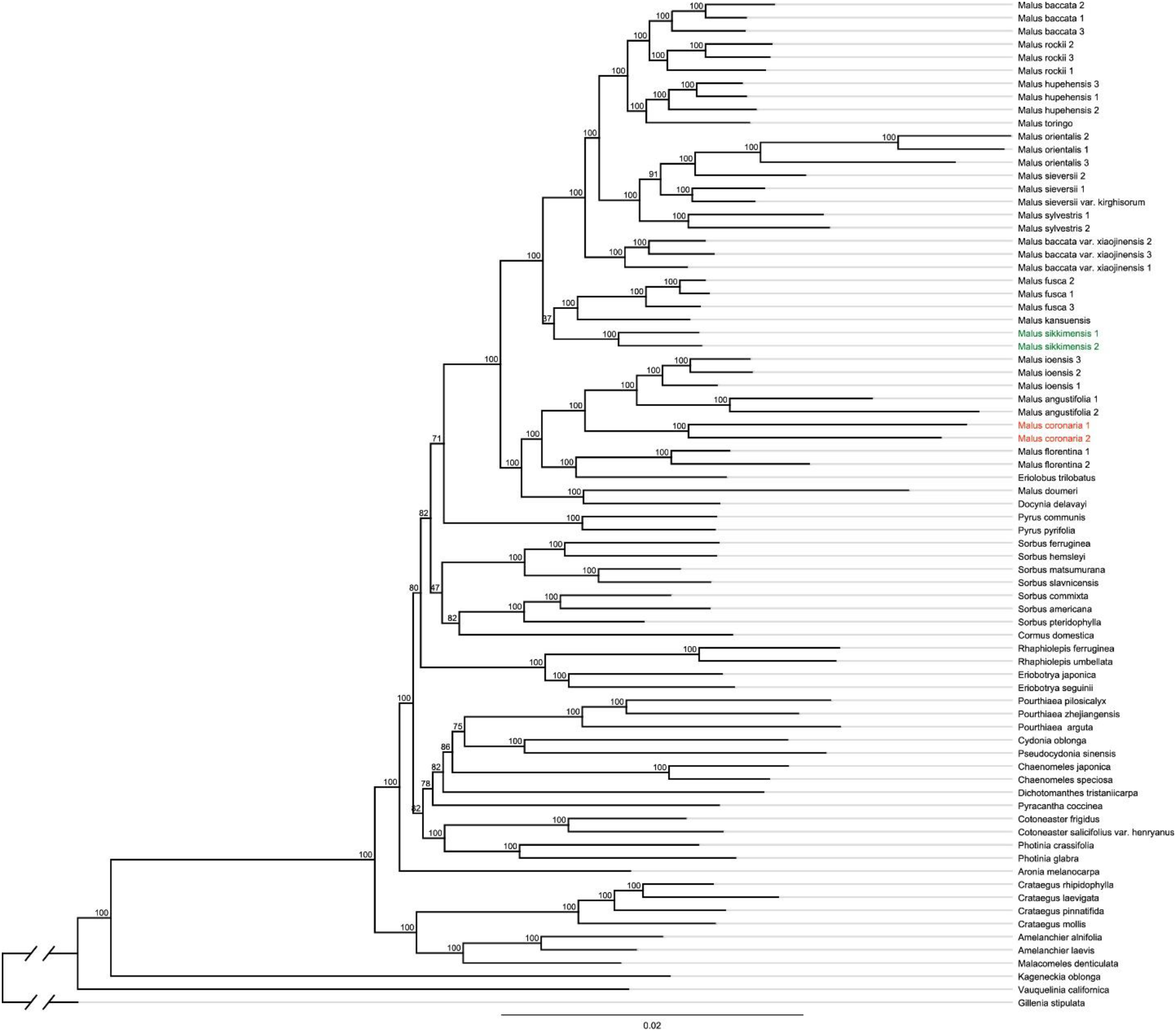
Maximum likelihood phylogeny of *Malus* s.l. in the framework of Maleae inferred from RAxML analysis of the concatenated 50%-sample dataset. Numbers above the branches indicate the bootstrap support.

**Fig. S2.**
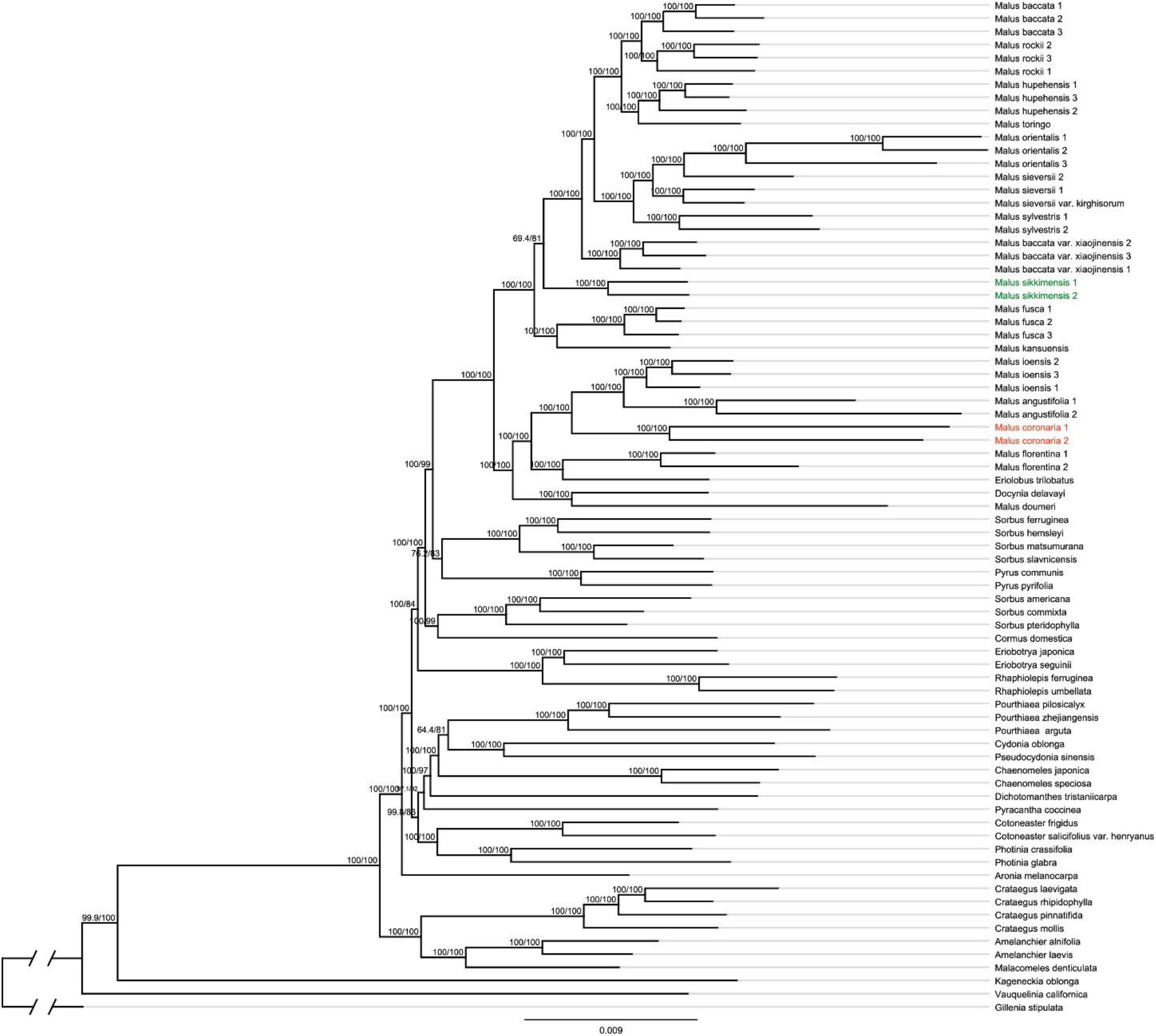
Maximum likelihood phylogeny of *Malus* s.l. in the framework of Maleae inferred from IQ-TREE2 analysis of the concatenated 50%-sample dataset. Numbers above the branches indicate the SH-aLRT support and Ultrafast Bootstrap support.

**Fig. S3.**
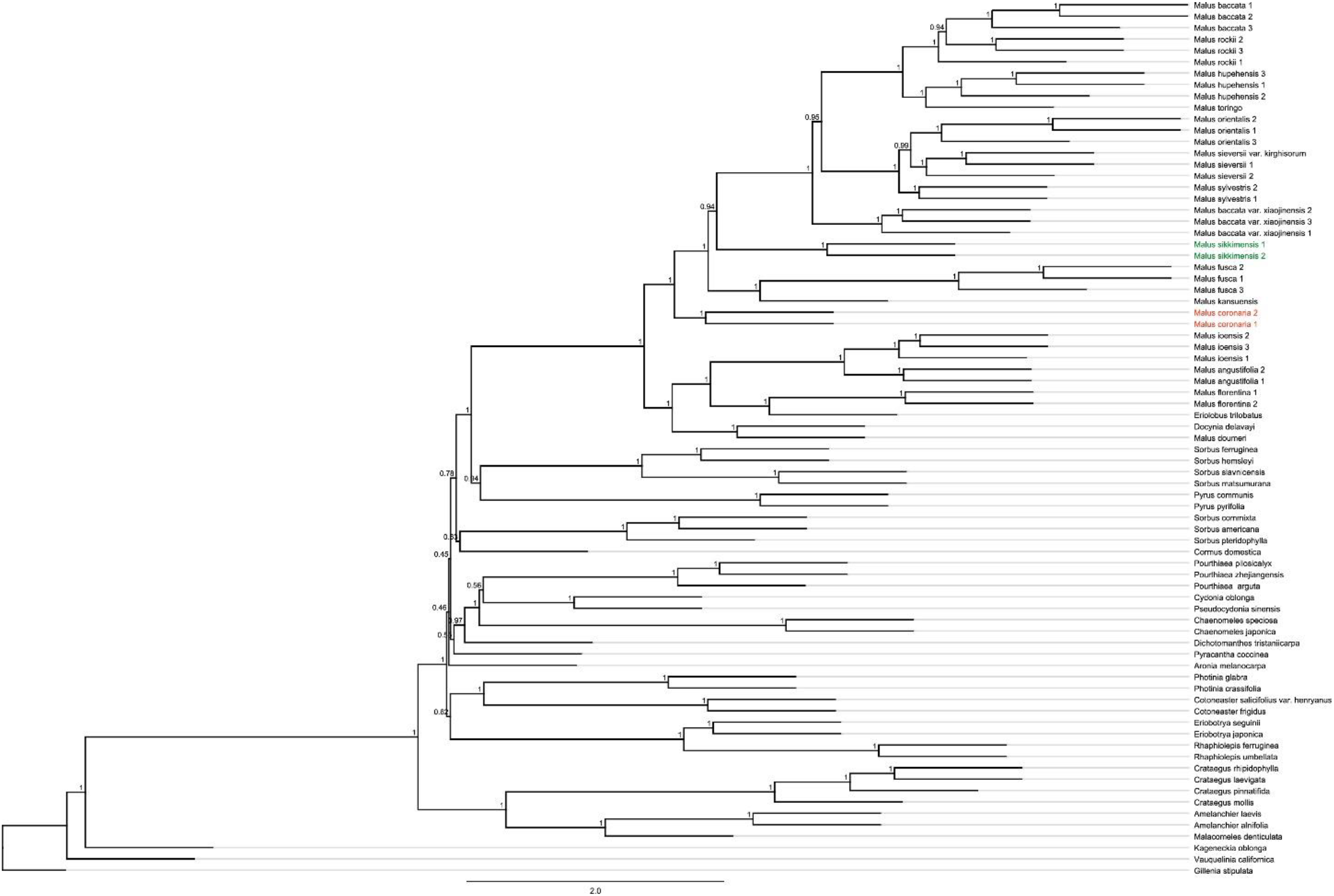
Species tree of *Malus* s.l. in the framework of Maleae inferred from ASTRAL-III of the concatenated 50%-sample dataset. Numbers above the branches indicate the branch support values measuring the support for a local posterior possibility.

**Fig. S4.**
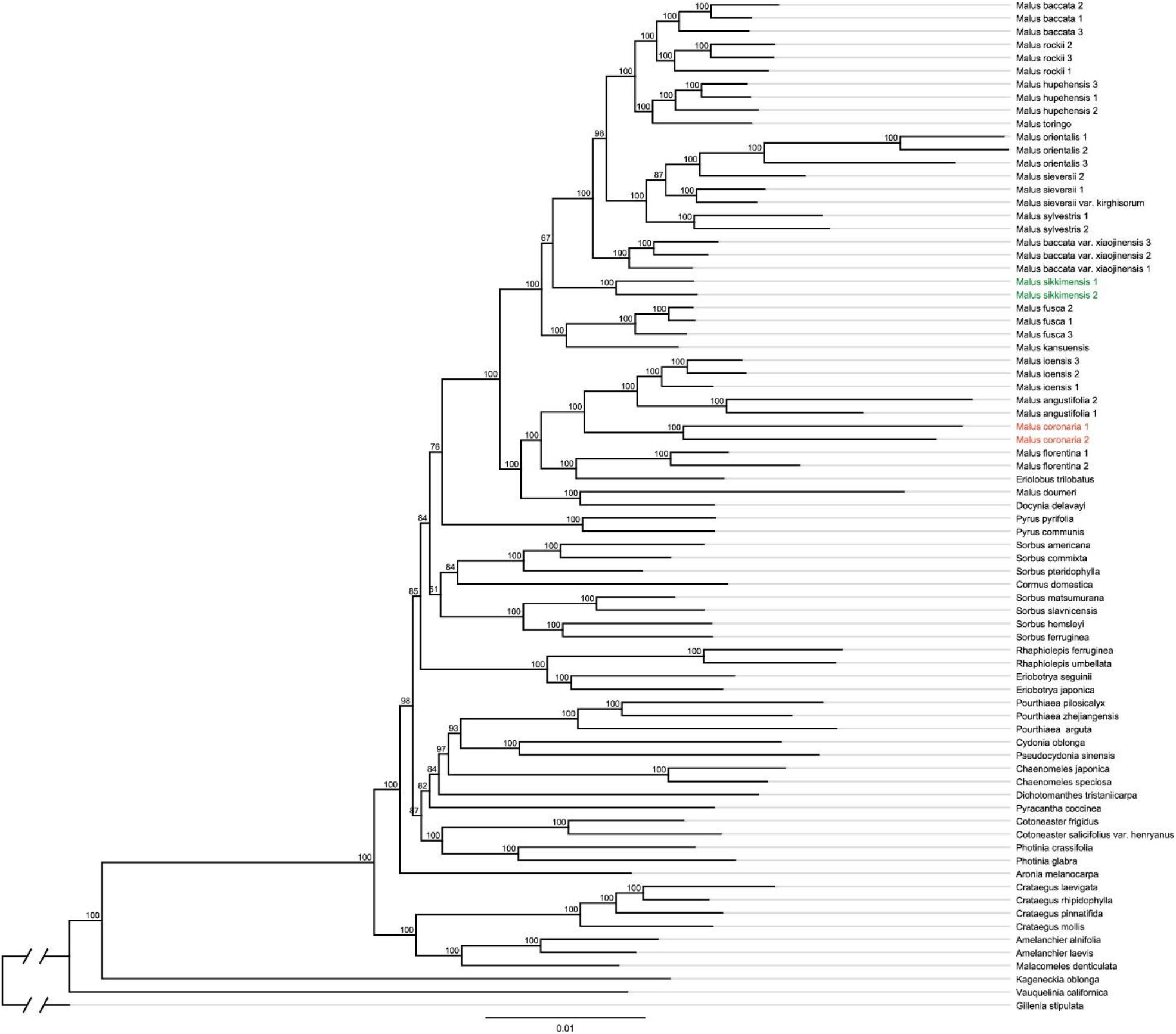
Maximum likelihood phylogeny of *Malus* s.l. in the framework of Maleae inferred from RAxML analysis of the concatenated 80%-sample dataset. Numbers above the branches indicate the bootstrap support (BS).

**Fig. S5.**
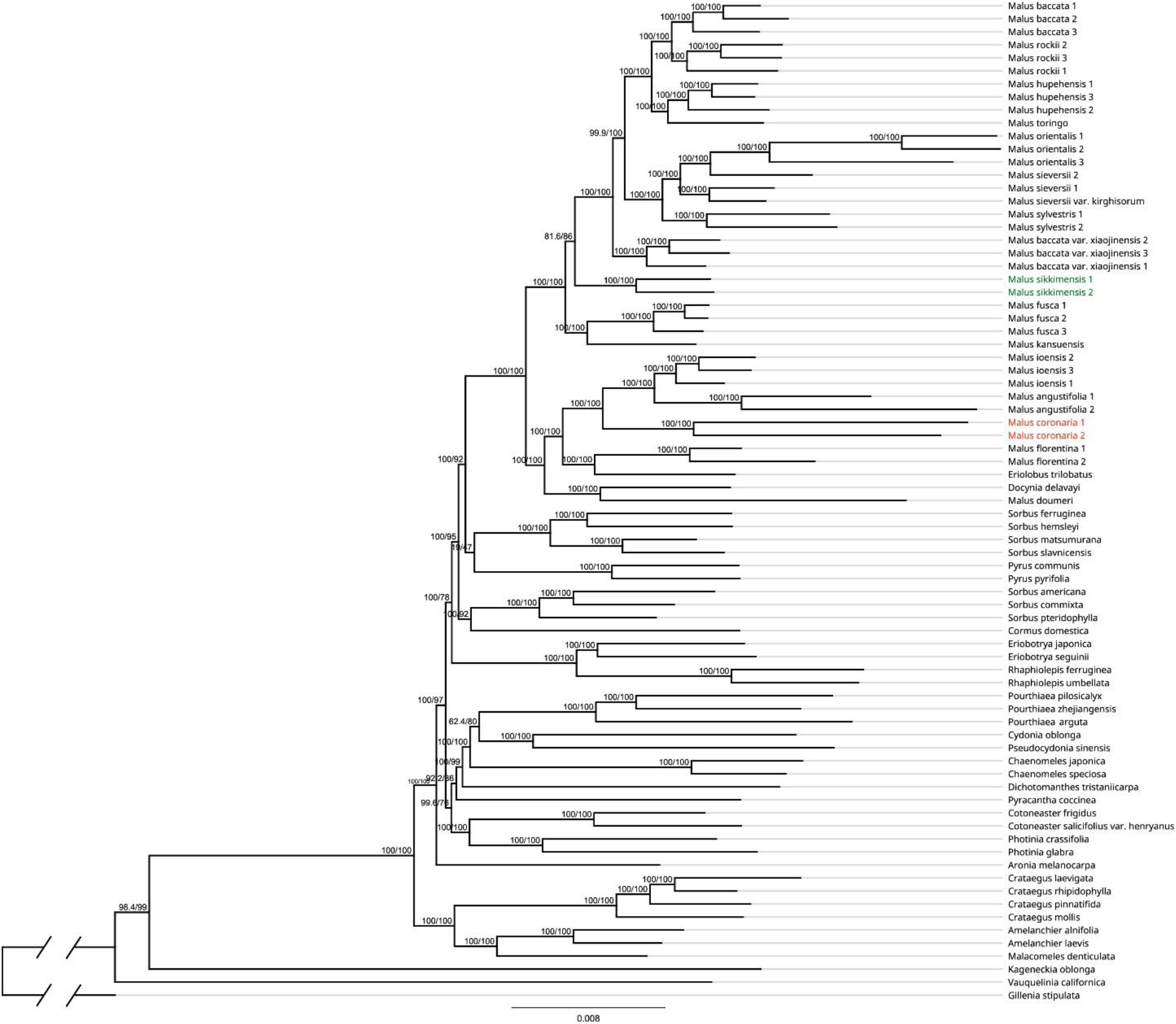
Maximum likelihood phylogeny of *Malus* s.l. in the framework of Maleae inferred from IQ-TREE2 analysis of the concatenated 80%-sample dataset. Numbers above the branches indicate the SH-aLRT support and Ultrafast Bootstrap (UFBoot) support.

**Fig. S6.**
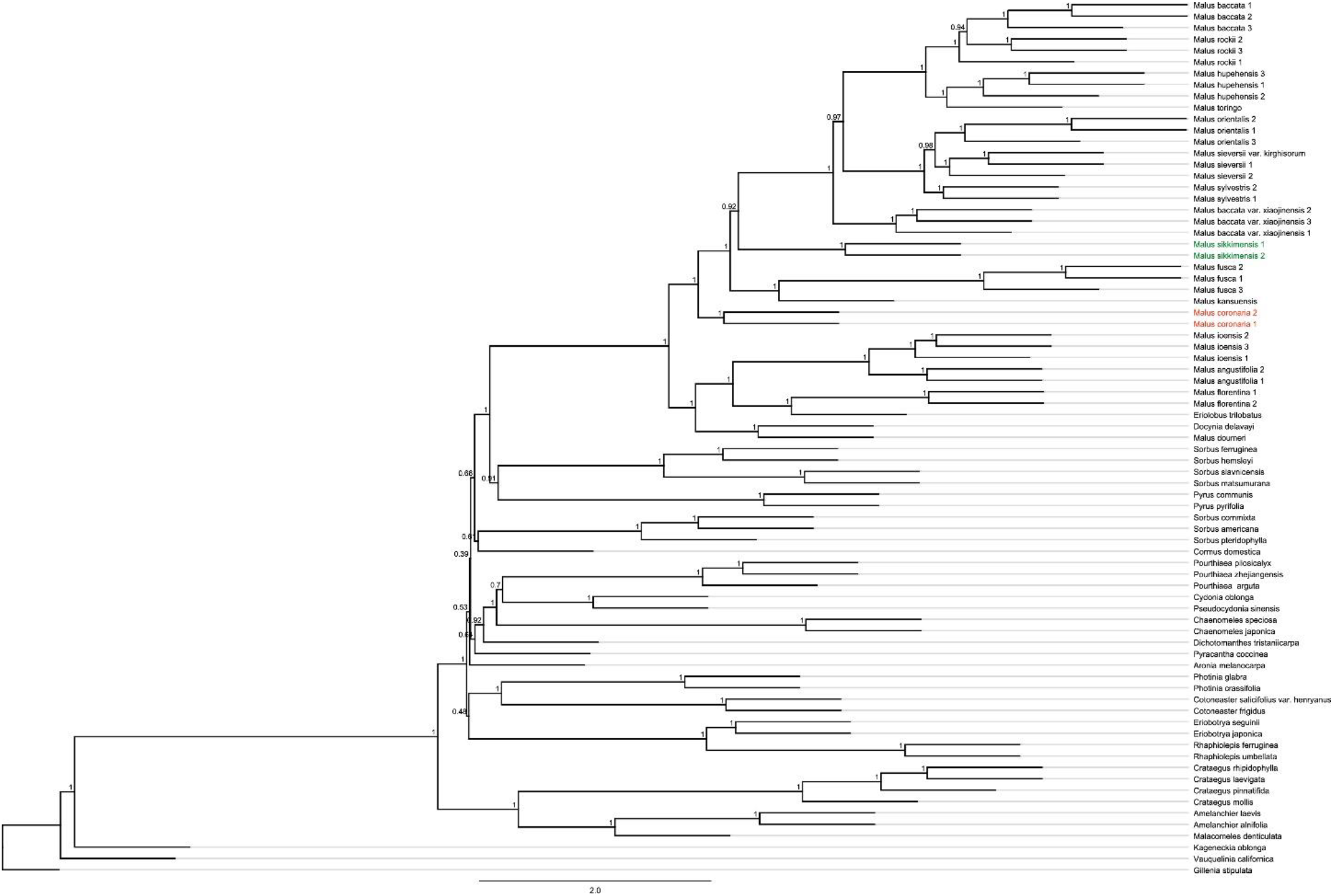
Species tree of *Malus* s.l. in the framework of Maleae inferred from ASTRAL-III of the concatenated 50%-sample dataset. Numbers above the branches indicate the branch support values measuring the support for a local posterior possibility.

**Fig. S7.**
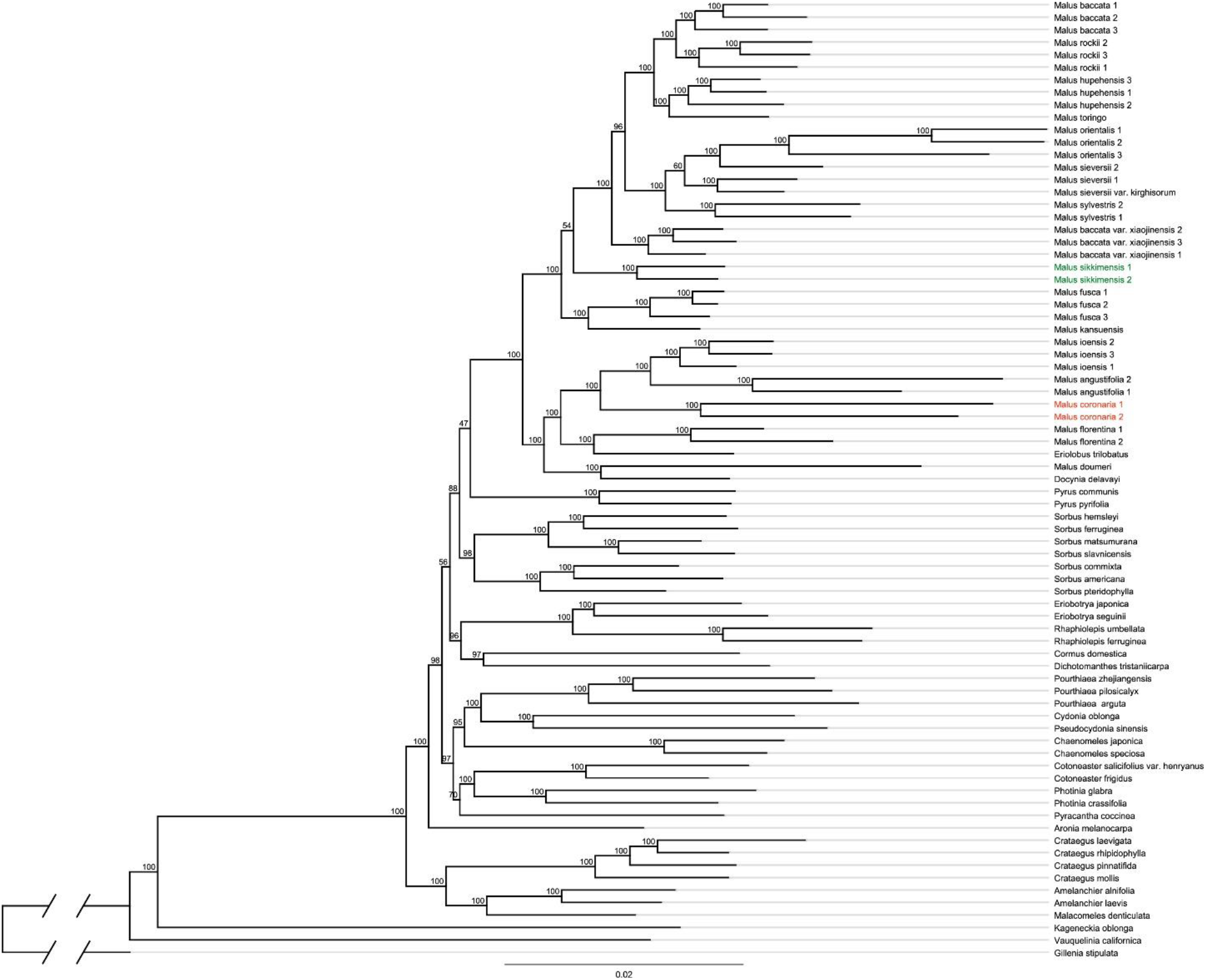
Maximum likelihood phylogeny of *Malus* s.l. in the framework of Maleae inferred from RAxML analysis of the all-sample dataset. Numbers above the branches indicate the bootstrap support (BS).

**Fig. S8.**
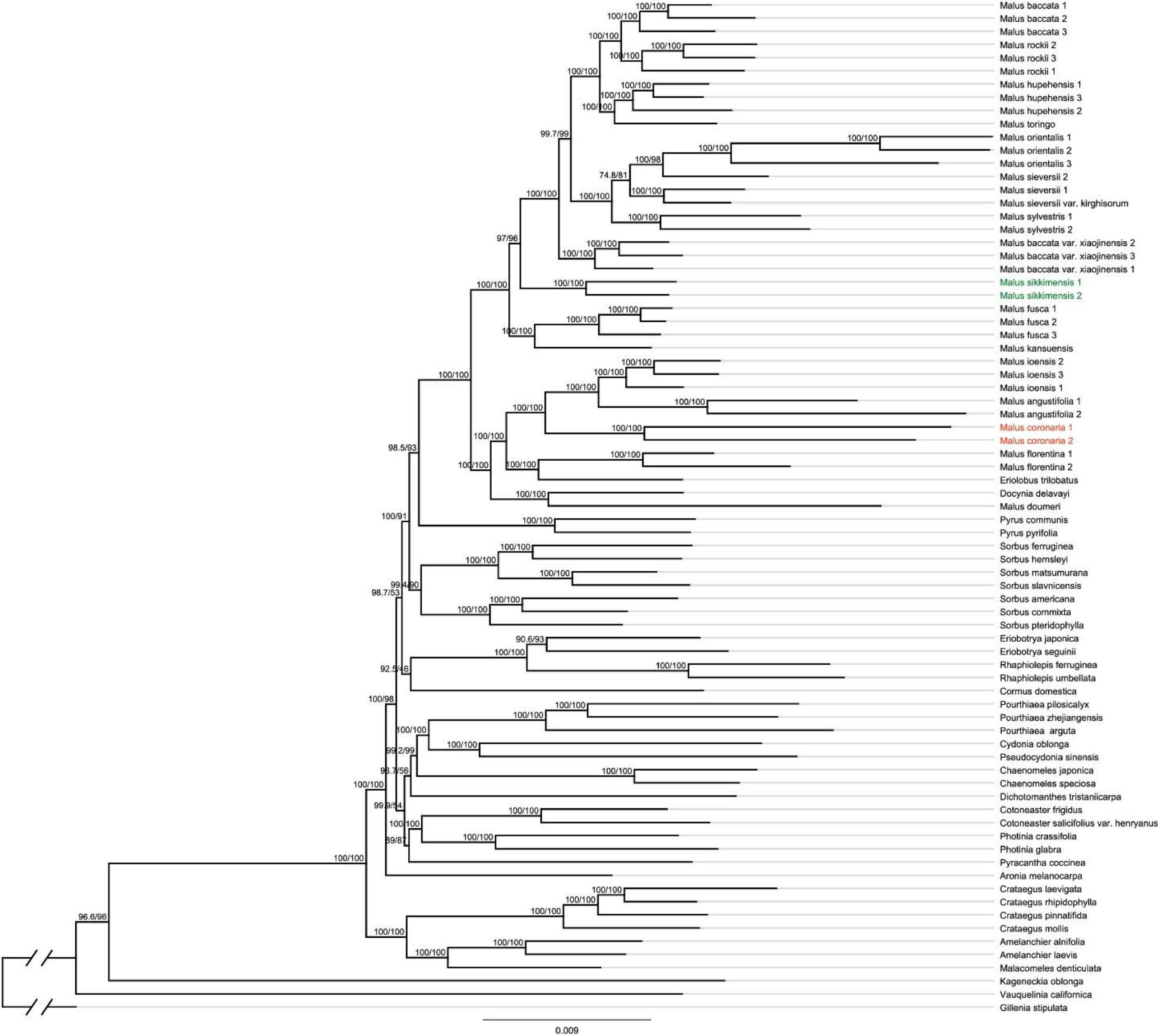
Maximum likelihood phylogeny of *Malus* s.l. in the framework of Maleae inferred from IQ-TREE2 analysis of the all-sample dataset. Numbers above the branches indicate the SH-aLRT support and Ultrafast Bootstrap (UFBoot) support.

**Fig. S9.**
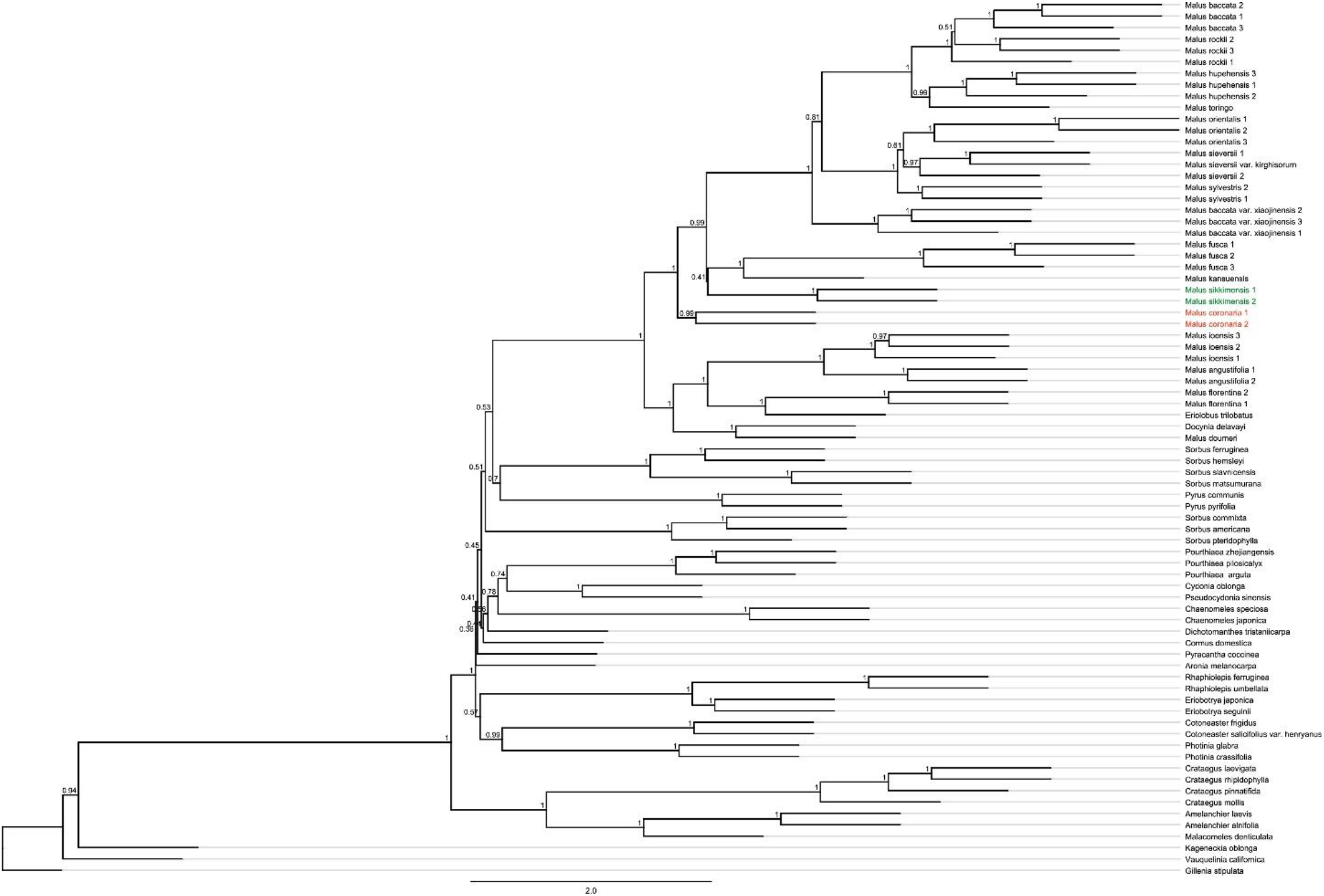
Species tree of *Malus* s.l. in the framework of Maleae inferred from ASTRAL-III of the all-sample dataset. Numbers above the branches indicate the branch support values measuring the support for a local posterior possibility.

**Fig. S10.**
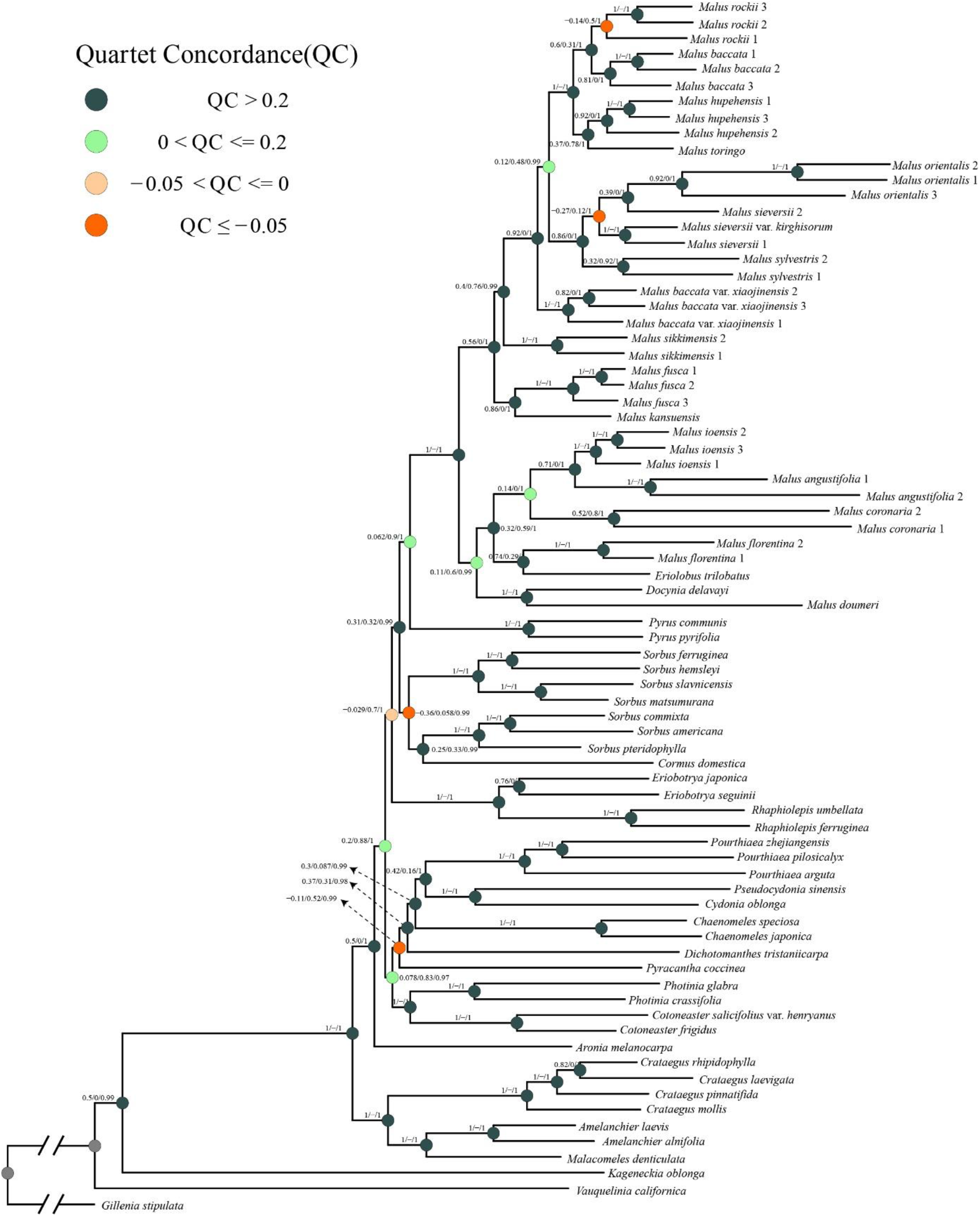
Maximum likelihood phylogeny of *Malus* s.l. in the framework of Maleae inferred from RAxML analysis of the concatenated 80%-sample dataset. Quartet Samping Scores are shown on branches indicating Quartet Concordance/Quartet Differential/Quartet Informativeness. Quartet Concordance is color-coded according to the legend.

**Fig. S11.**
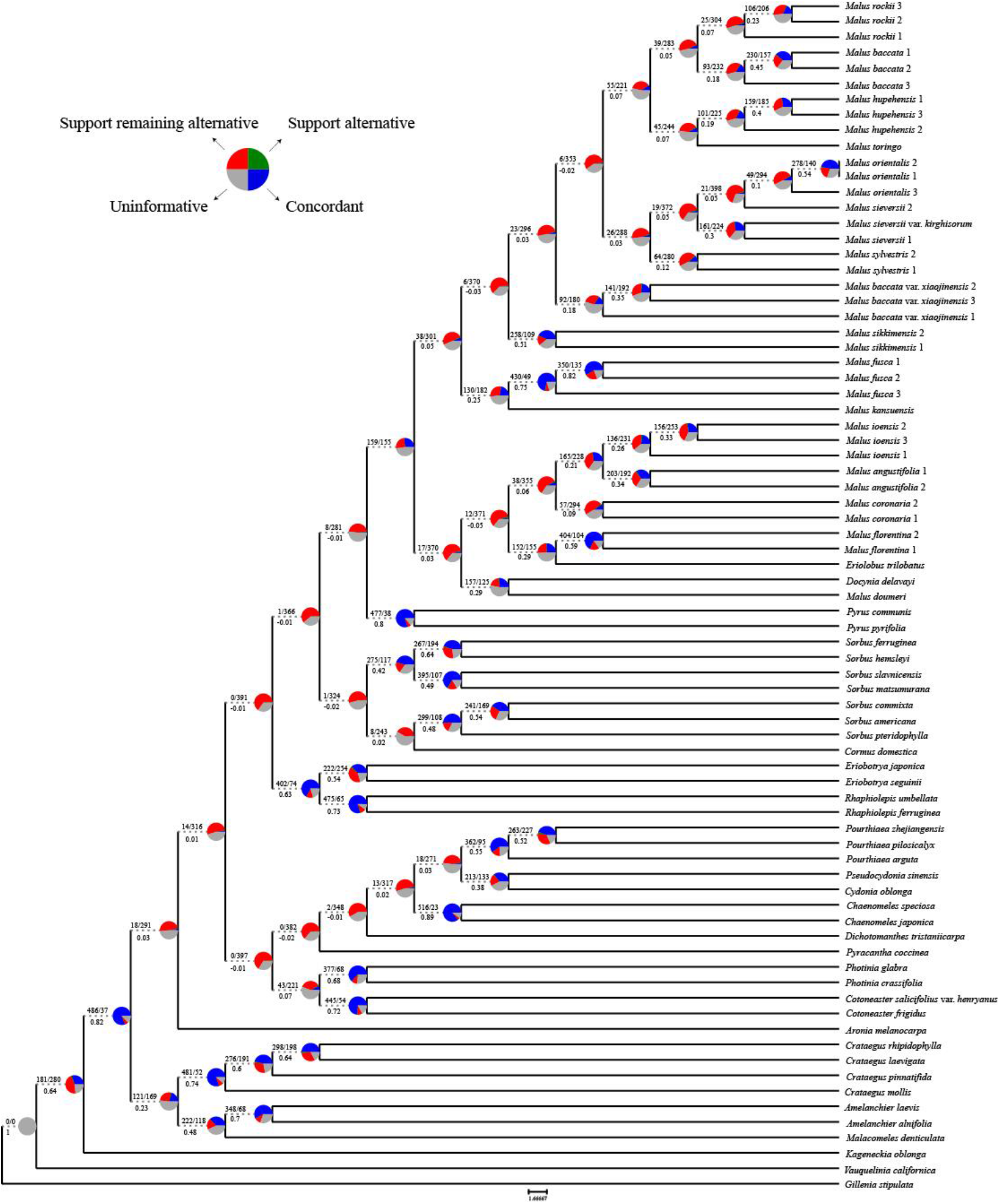
Maximum likelihood cladogram of *Malus* s.l. inferred from RAxML analysis of the concatenated 80%-sample dataset. Numbers above branches indicate the number of gene trees concordant/conflicting with that node in the species tree. Numbers below the branches are the Internode Certainty All (ICA) score. Pie charts on nodes present the proportion of gene trees that support that clade (blue), the proportion that support the main alternative bifurcation (green), the proportion that support the remaining alternatives (red), and the proportion (conflict or support) that have < 50% bootstrap support (gray).

**Fig. S12.**
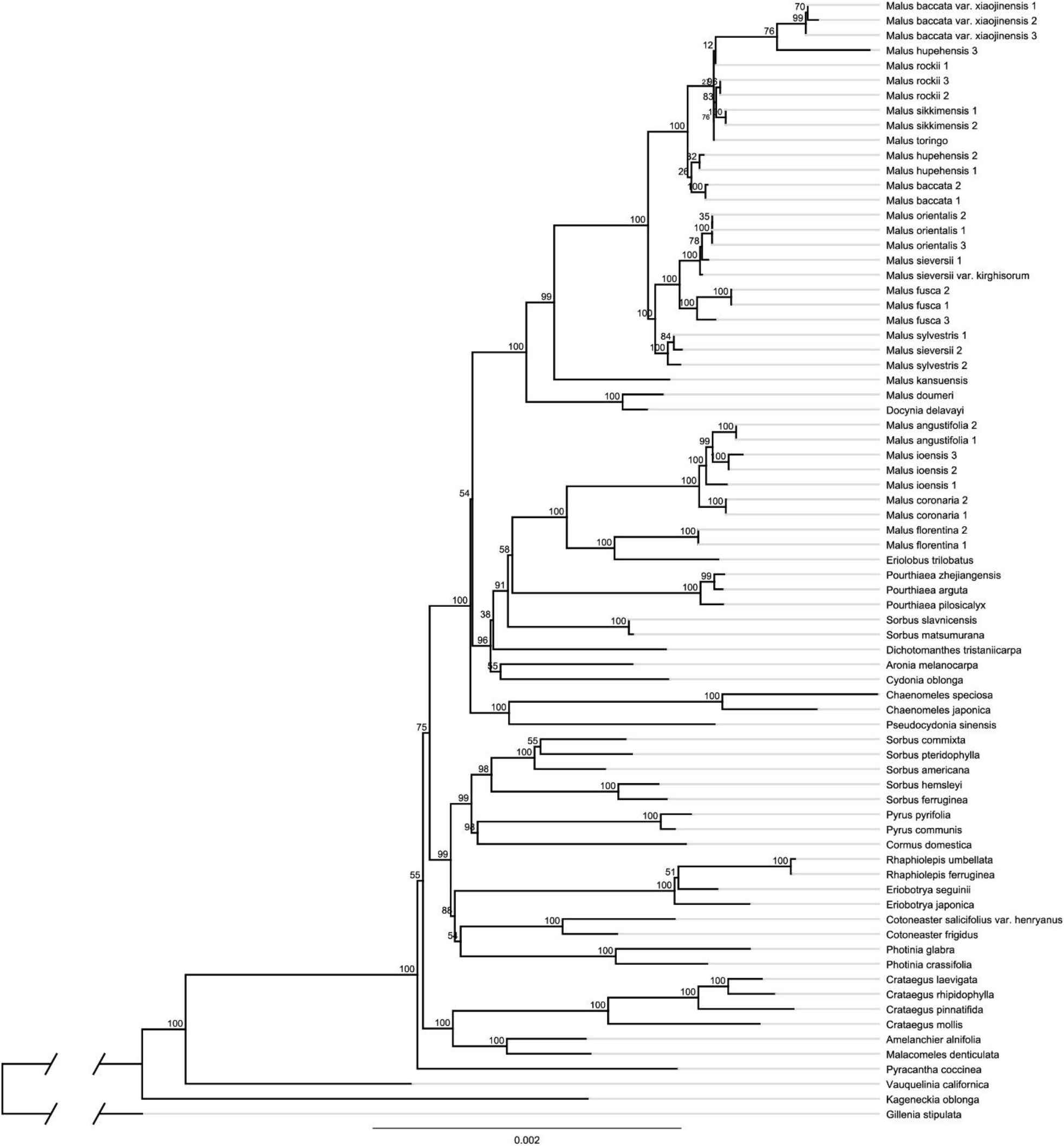
Maximum likelihood phylogeny of *Malus* s.l. in the framework of Maleae inferred from RAxML analysis of the concatenated 80 plastid coding genes. Numbers above the branches indicate the bootstrap support.

**Fig. S13.**
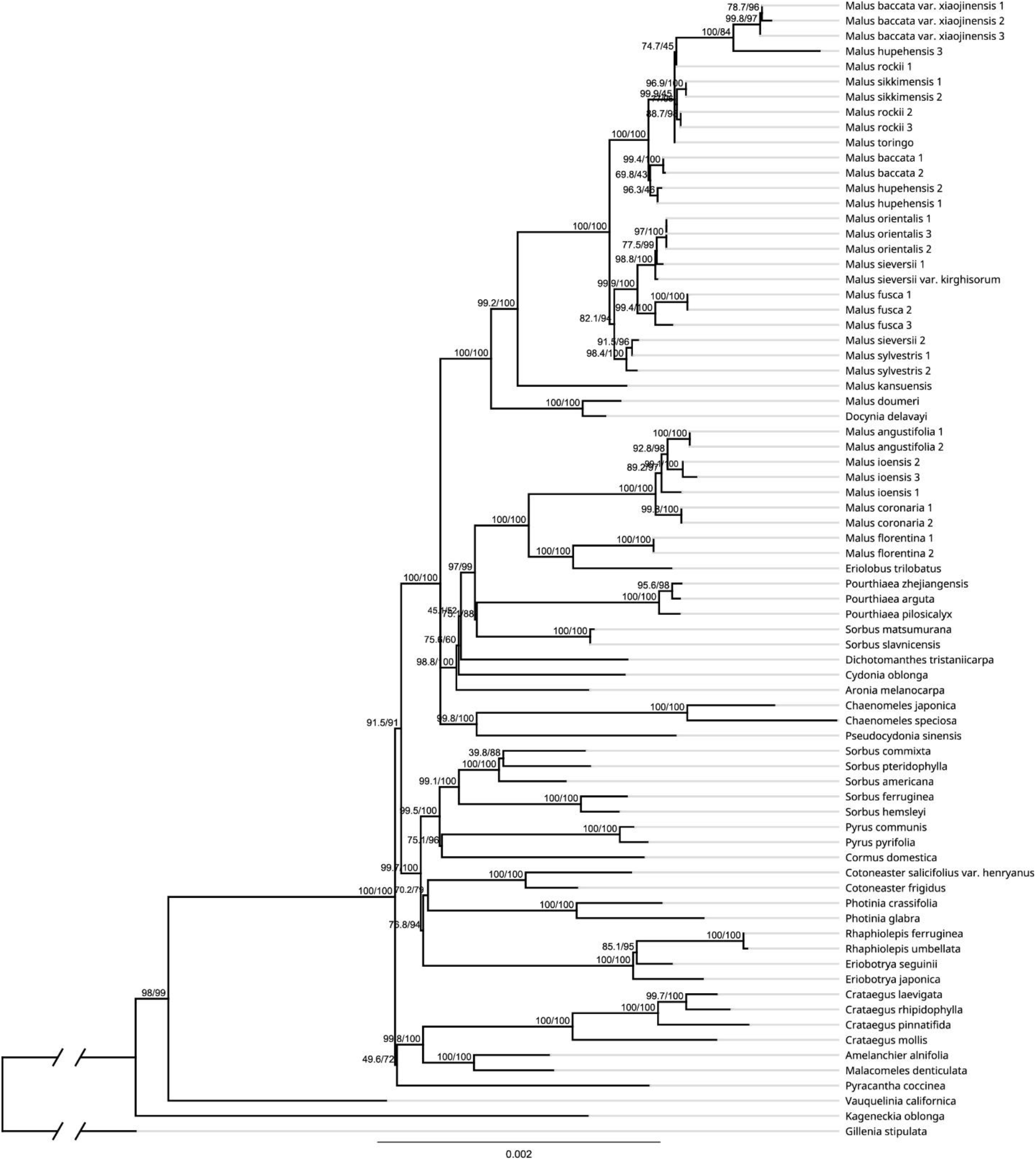
Maximum likelihood phylogeny of *Malus* s.l. in the framework of Maleae inferred from IQ-TREE2 analysis of the concatenated 80 plastid coding genes. Numbers above the branches indicate the SH-aLRT support and Ultrafast Bootstrap support.

**Fig. S14.**
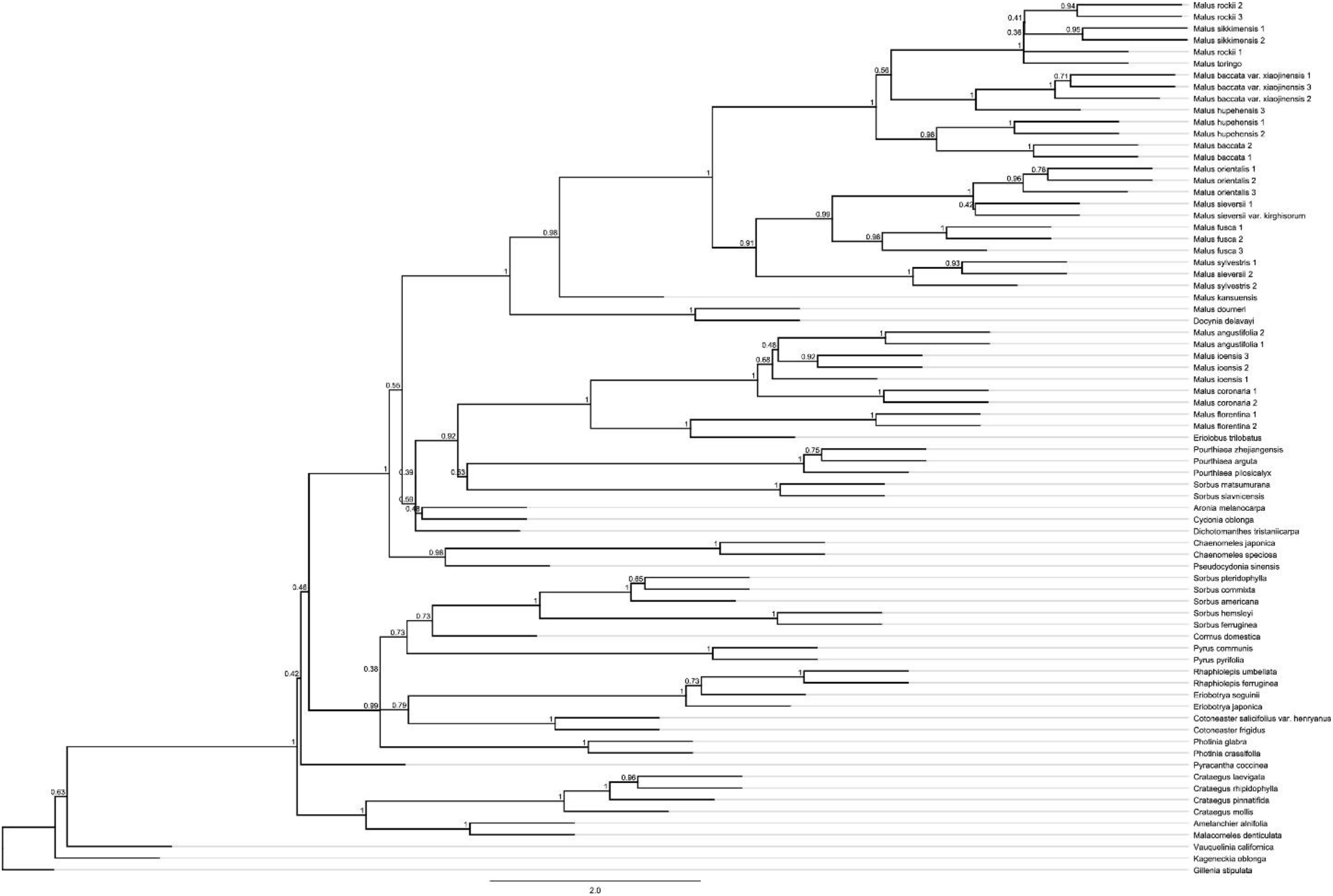
Species tree of *Malus* s.l. in the framework of Maleae inferred from ASTRAL-III of the concatenated 80 plastid coding genes. Numbers above the branches indicate the branch support values measuring the support for a local posterior possibility.

**Fig. S15.**
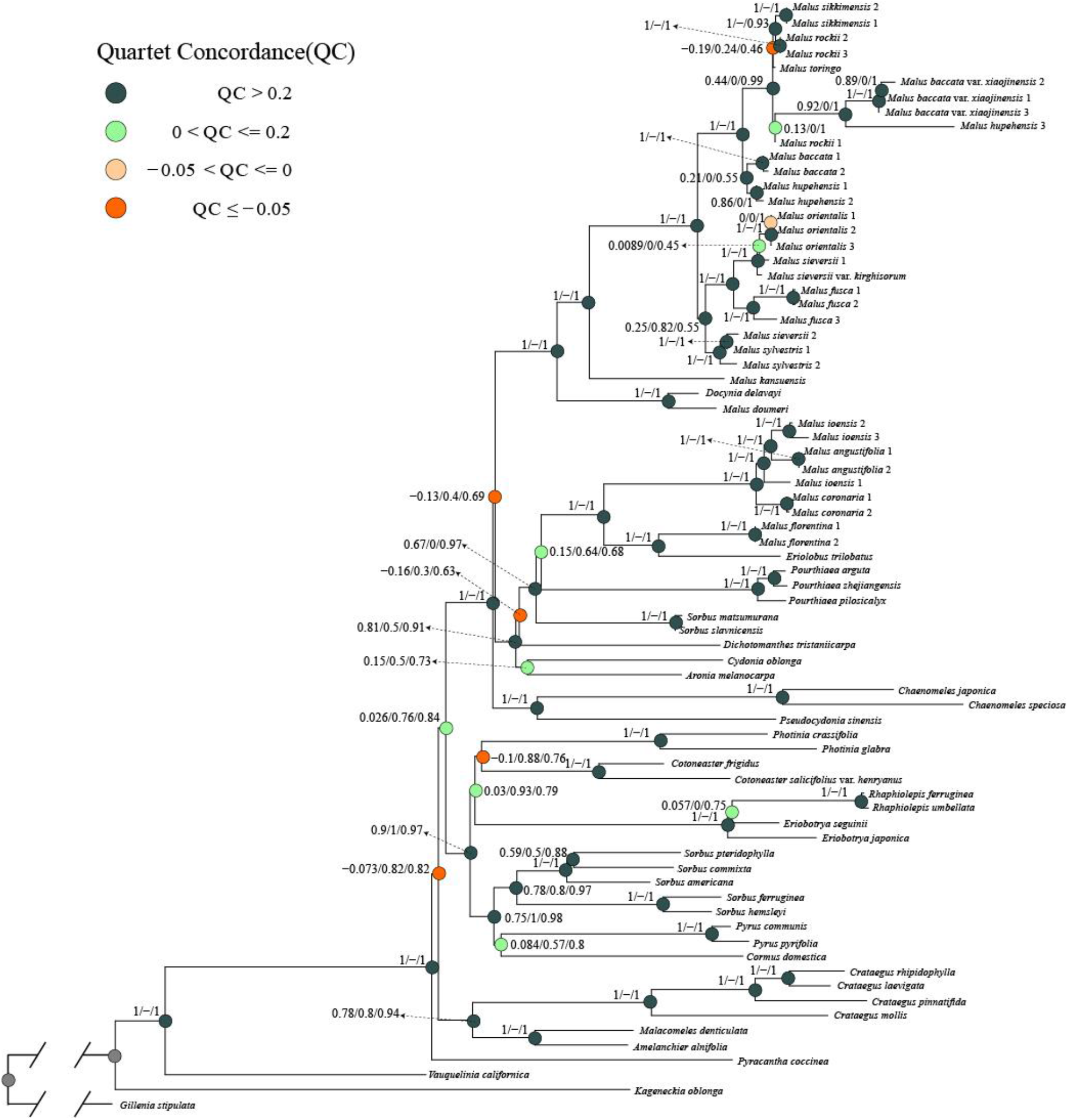
Maximum likelihood phylogeny of *Malus* s.l. in the framework of Maleae inferred from RAxML analysis of the concatenated 80 plastid coding genes. Quartet Samping Scores are shown on branches indicating Quartet Concordance/Quartet Differential/Quartet Informativeness. Quartet Concordance is color-coded according to the legend.

**Fig. S16.**
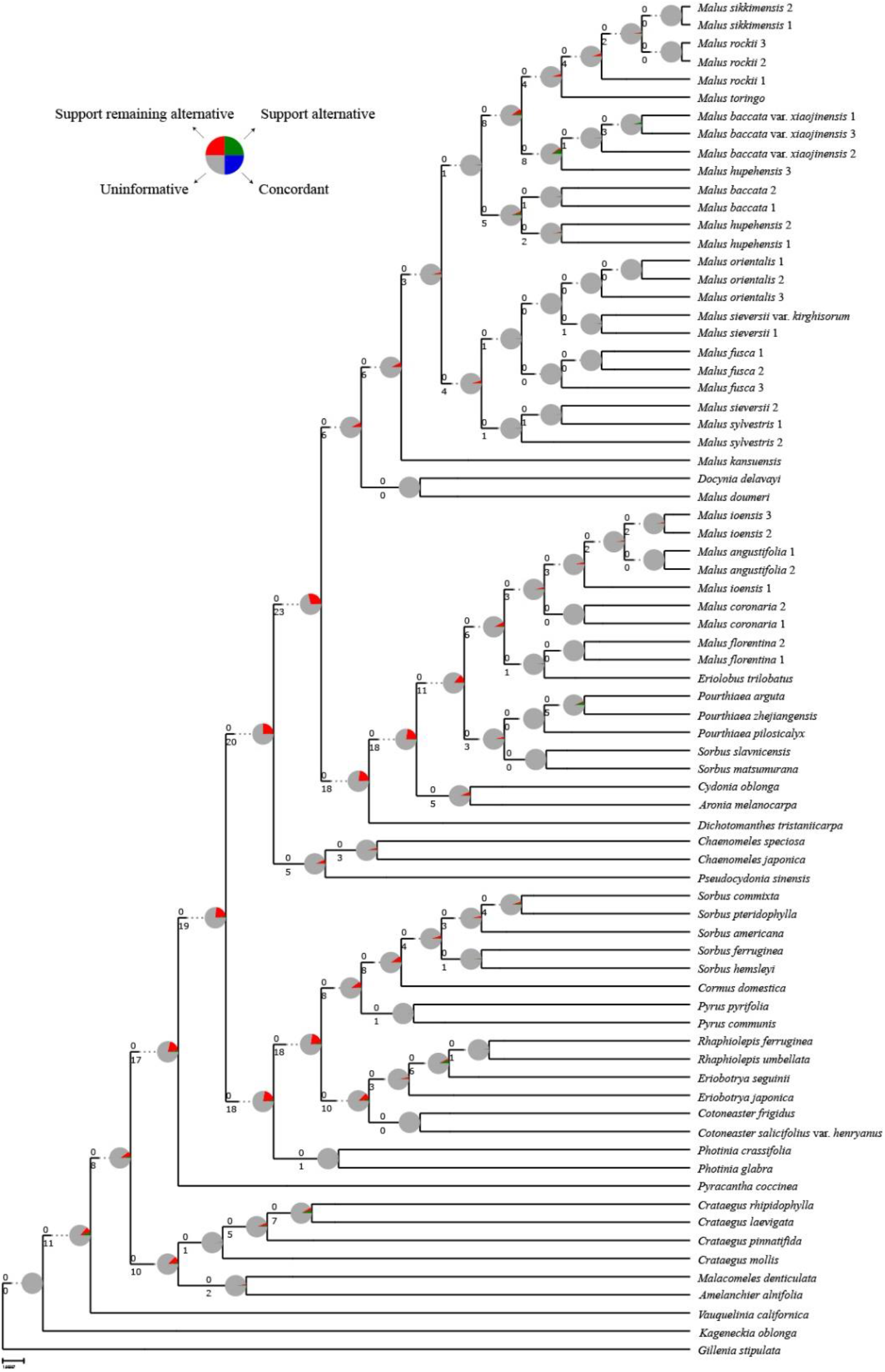
Maximum likelihood cladogram of *Malus* s.l. inferred from RAxML analysis of the concatenated 80 plastid coding genes. Numbers above branches indicate the number of gene trees concordant/conflicting with that node in the species tree. Numbers below the branches are the Internode Certainty All (ICA) score. Pie charts on nodes present the proportion of gene trees that support that clade (blue), the proportion that support the main alternative bifurcation (green), the proportion that support the remaining alternatives (red), and the proportion (conflict or support) that have < 50% bootstrap support (gray).

**Fig. S17.**
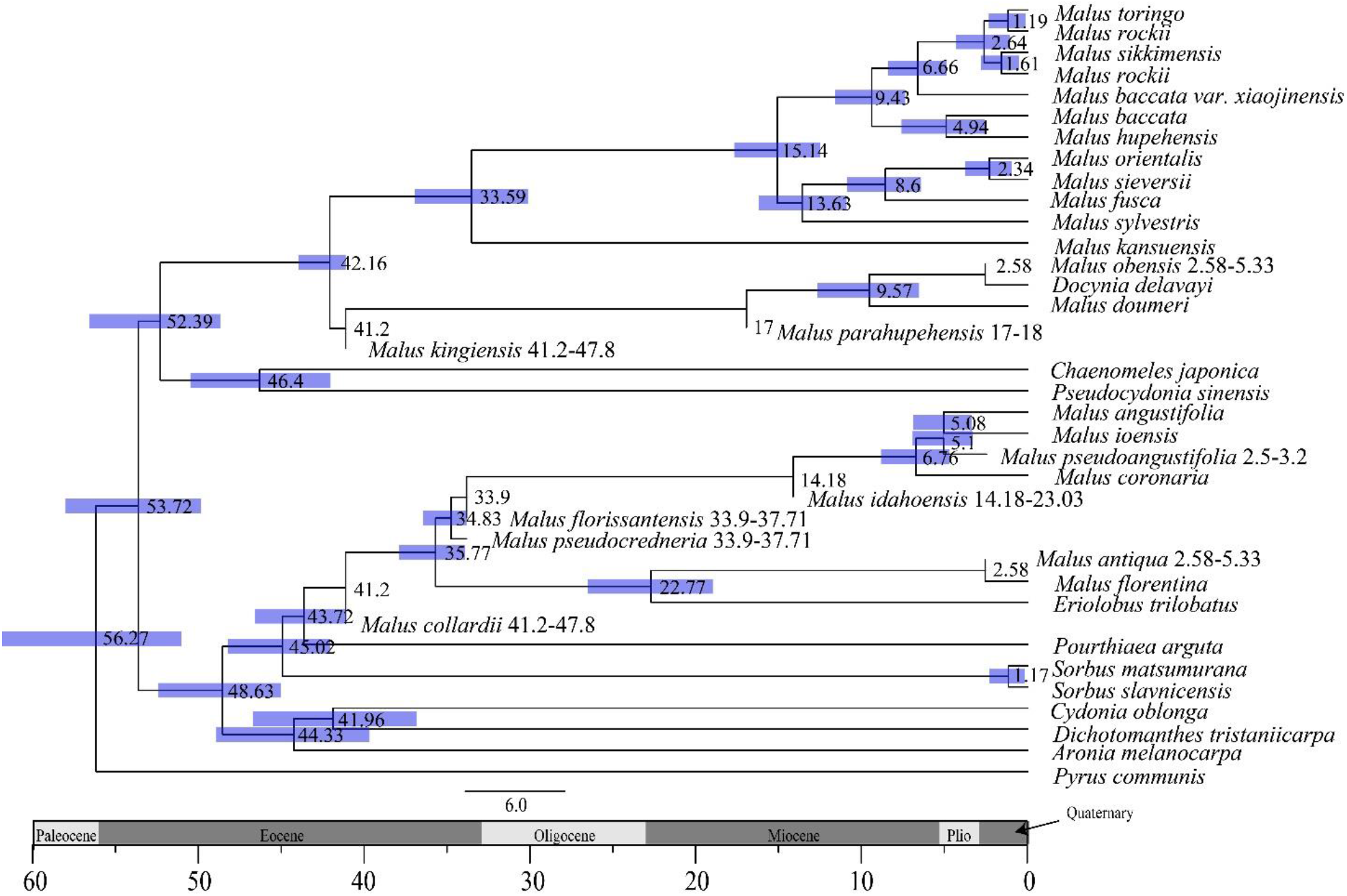
Dated chronogram for the apple genus *Malus* inferred from BEAST with the Fossilized Birth-Death process based on the 80 plastid codinig genes dataset.

**Fig. S18.**
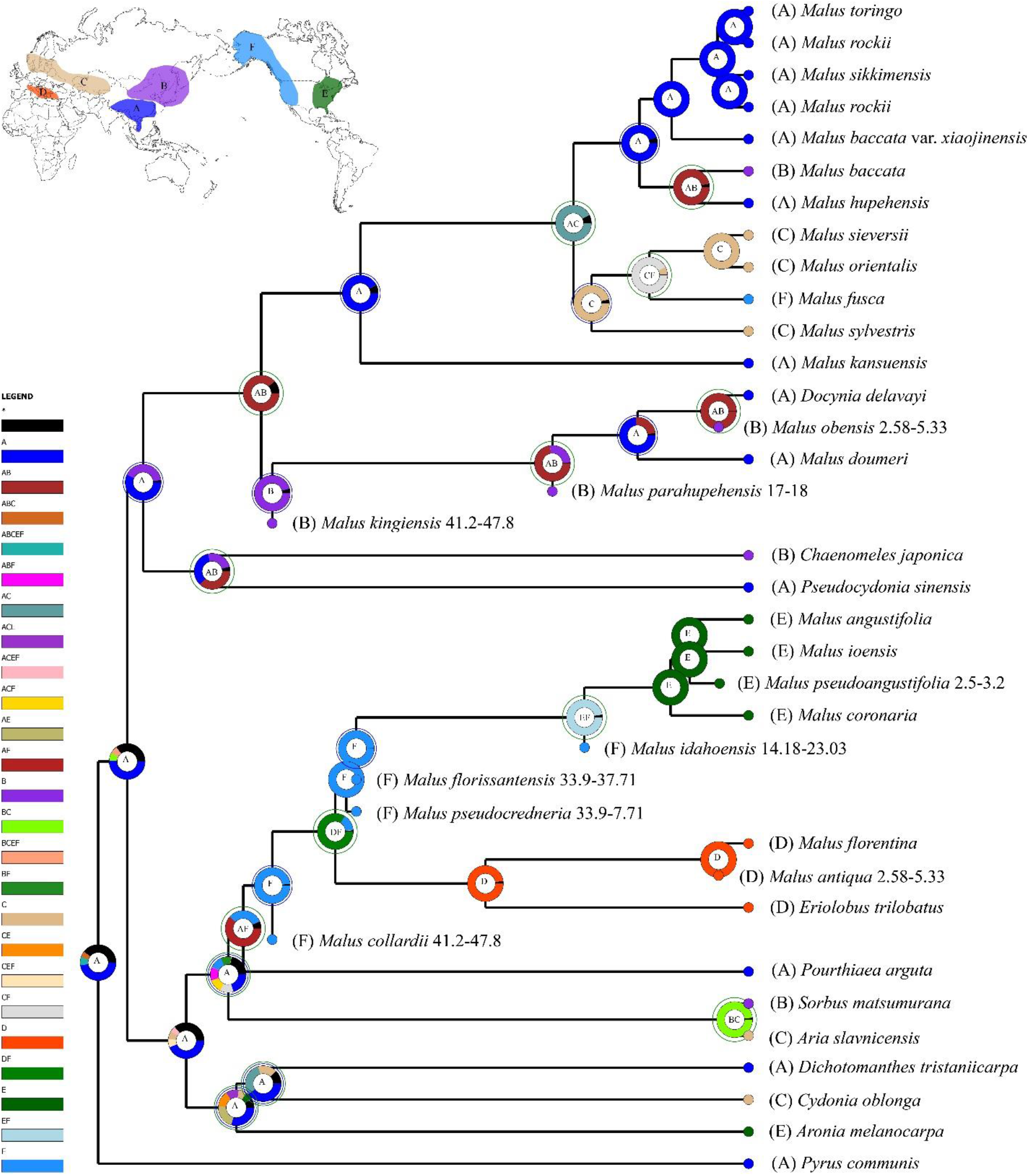
The ancestral area reconstruction using BioGeoBEARS from the 80 plastid coding genes dataset, with the colored key identifying extant and possible ancestral ranges, (A), Southern East Asia; (B), Northern East Asia; (C), Europe and Central Asia; (D), Mediterranean; (E), Eastern North America; (F), Western North America.

